# Vein fate determined by flow-based but time-delayed integration of network architecture

**DOI:** 10.1101/2021.12.29.474405

**Authors:** Sophie Marbach, Noah Ziethen, Leonie Bastin, Felix K. Bäuerle, Karen Alim

**Author notes:** These two authors contributed equally (Sophie Marbach, Noah Ziethen).

## Abstract

Veins in vascular networks, such as in blood vasculature or leaf networks, continuously reorganize, grow or shrink, to minimize energy dissipation. Flow shear stress on vein walls has been set forth as the local driver for a vein’s continuous adaptation. Yet, shear feedback alone cannot account for the observed diversity of vein dynamics – a puzzle made harder by scarce spatiotemporal data. Here, we resolve network-wide vein dynamics and shear rate during spontaneous reorganization in the prototypical vascular networks of *Physarum polycephalum*. Our experiments reveal a plethora of vein dynamics (stable, growing, shrinking) where the role of shear is ambiguous. Quantitative analysis of our data reveals that (a) shear rate indeed feeds back on vein radius, yet, with a time delay of 1 — 3 min. Further, we reconcile the experimentally observed disparate vein fates by developing a model for vein adaptation within a network and accounting for the observed time delay. The model reveals that (b) vein fate is determined by parameters – local pressure or relative vein resistance – which integrate the entire network’s architecture, as they result from global conservation of fluid volume. Finally, we observe avalanches of network reorganization events that cause entire clusters of veins to vanish. Such avalanches are consistent with network architecture integrating parameters governing vein fate as vein connections continuously change. As the network architecture integrating parameters intrinsically arise from laminar fluid flow in veins, we expect our findings to play a role across flow-based vascular networks.

---

Veins interwebbed in networks distribute resources across numerous forms of life, from the blood vasculature in animals [1–4], via the leaf venation in plants [5, 6] to the vein networks entirely making up fungi and slime molds [7, 8]. Continuous reorganization is integral to a network’s success: veins perpetually grow and shrink [3, 9, 10]. While vein dynamics are usually observed for individual veins [1], reorganization patterns at the network scale remain a puzzle. Yet, understanding network reorganization is crucial to shed light on the mechanics of development [3] and widespread diseases [11, 12].

While the biological makeup of vasculature systems is quite diverse, the physics that governs pervading and laminar fluid flows is the same [13]. Already almost a century ago, Murray introduced the idea that shear stress exerted by fluid flows on a vein wall determines vein radius size [14]. Within his framework, at steady state, veins minimize viscous dissipation while constrained by a constant metabolic cost to sustain the vein. Solving the minimization problem yields that shear stress, driver of viscous dissipation, should be constant among veins. Since Murray derived his hypothesis, studies have focused on *static* networks [6, 15, 16]. Data on optimal static network morphologies agrees very well with Murray’s predictions, strikingly across very different forms of life; from animals [17, 18], to plants [17, 19] and slime molds [20, 21]. Fluid flow physics is, therefore, key to understanding vascular morphologies.

Beyond steady state, during reorganization, how do flows shape network morphologies? Data on vein *dynamics*, [3, 22–25], even during spontaneous reorganization, is limited due to the difficulty of acquiring time-resolved data covering entire networks. Observation of network excerpts suggests that flow shear stress alone can not account for the diversity of observed dynamics [26]. In light of scarce experimental observations, a number of vein adaptation models have been introduced [4, 10, 20, 22, 27–33]. Yet, the mechanisms that govern vein adaptation and thereby network reorganization can only be conclusively determined experimentally.

Here, we investigate the vascular networks formed by the slime mold *Physarum polycephalum*. Since the organisms’ body is reduced to approximately two dimensions [8, 21, 22], it opens up the unique possibility to quantify vein dynamics and fluid flows simultaneously in the entire network. From the fluid flows, we then quantify shear rate, directly related to shear stress by the inverse of the fluid’s dynamic viscosity. Flows in the veins arise from rhythmic contractions of vein walls due to actomyosin activity in the vein cortex. As the flows oscillatory component changes rapidly on 1 min to 2 min [34, 35], average flows dominate long-term vein adaptation dynamics on 10 min and more. Our aim, here, is to employ *P. polycephalum* to quantify experimentally and rationalize *individual* and *global* vein reorganization dynamics.

Our quantitative data reveals that shear rate indeed feeds back on vein radii, notably with a time delay. Furthermore, the effect of shear rate is disparate: similar shear rate values may cause veins either to grow or to shrink. To reconcile these disparate dynamics, we derive a model of vein adaptation in networks based on Kirchhoff’s laws. Our model reproduces experimental observations and predicts that shear rate is not the only driver of vein adaptation, but also network-integrating parameters take control: fluid pressure and relative vein resistance. Both parameters integrate the network’s architecture since they derive from fluid volume conservation on the network scale expressed by Kirchhoff’s laws. As veins shrink and grow, network architecture continuously changes. As a consequence, a vein’s fate to remain or shrink, is not predetermined by the current static network architecture but rather changes in time. This dynamic perspective explains avalanches of shrinking and disappearing veins in connected clusters. The mechanistic insight gained by our model suggests that the rules of vein reorganization, particularly the role of network-integrating parameters like fluid pressure and relative vein resistance, might be critical to understanding vascular networks across different life forms.

## I. INDIVIDUAL VEIN DYNAMICS HAVE COMPLEX SHEAR RATE-RADIUS RELATION

### A. Quantifying vein dynamics

We observe vein dynamics in *P. polycephalum* specimen using two complementary imaging techniques, either close-up observation of single veins or full network imaging (Fig. 1 and additional methods in Appendix 1). Close-up vein microscopy over long timescales (Fig. 1-A.i, see also Movie 1) allows us to directly measure radius dynamics *a*(*t*) and velocity profiles *v*(*r,t*) inside vein segments using particle image velocimetry (Fig. 1-A.ii), where *t* is time and r is the radial coordinate along the tube. From velocity profiles, we extract the flow rate across a vein’s cross-section *Q*(*t*) = 2π ∫ *v*(*r,t*)*rdr*. In full networks (Fig. 1-B.i, see also Movie 2), radius dynamics *a(t*) are measured for each vein segment and flow rates *Q*(*t*) are subsequently calculated numerically integrating conservation of fluid volume via Kirchhoff laws, see Appendix 1.

**Figure 1.**
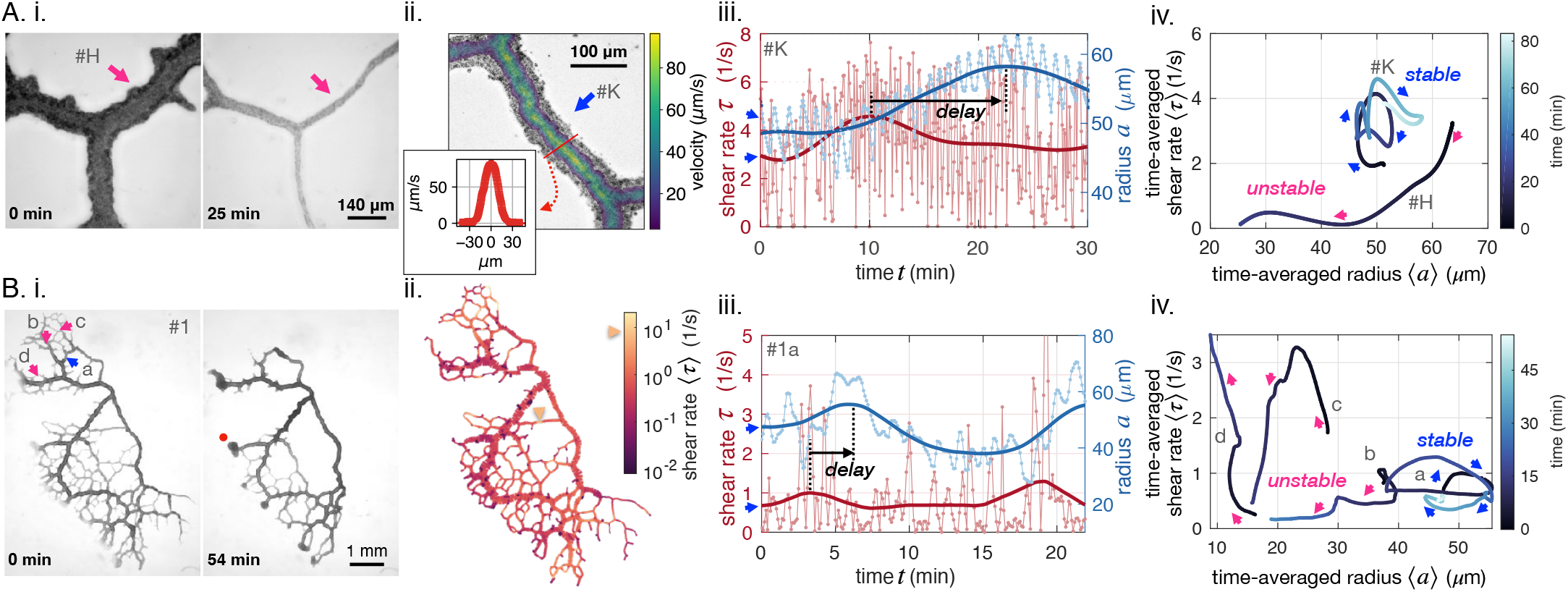
Diverse vein dynamics emerge during network reorganization. (A) Close-up and (B) full network analysis of vein radius dynamics and associated shear rate in *P. polycephalum*. (i) Bright-field images of reorganizing specimens allow us to record vein dynamics. (ii) Velocity measurements: (A) Velocity profiles along vein segments extracted with particle image velocimetry (inset: profile along vein cross-section) and (B) vein contractions driving internal flows over the entire network are integrated to calculate shear rate in veins (here shown at the initial observation time). The color scale indicates the magnitude of shear rate in each colored vein segment. For example, the yellow arrow points to a vein with a high calculated shear rate. (iii) Change in shear rate preceding changes in vein radius, both shown as a function of time (connected dots) and their time-averaged trends (full lines). A.iii shows the dynamics in the vein #*K* from A.ii, B.iii shows the vein marked in blue in B.i. (iv) The time-averaged shear rate versus the time-averaged radius displays circling dynamics for stable veins and diverse qualitative dynamics for unstable, vanishing veins. Blue color shades encode time. Trajectory arrow colors match arrow colors marking vein position in A.i (#*H*), A.ii (#*K*) and B.i, respectively. Veins marked in pink are shrinking, while stable veins are in blue.

Our imaging techniques resolve vein adaptation over a wide range of vein radii, *a* = 5 — 70 μm. Radii data show rhythmic peristaltic contractions, with a period of *T* ≃ 1 – 2 min (light blue in Fig. 1-iii). We calculate shear rate from fluid flows as 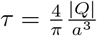. Unlike shear stress, shear rate measurements do not require knowledge of the fluid’s viscosity and are, therefore, more precise. Since both quantities are directly proportional, the conclusions we draw for shear rate apply to shear stress on the typical timescale of our experiments, where potential aging affects altering fluid viscosity can be neglected. We observe that shear rate *τ* oscillates with twice the contraction frequency (light red in Fig. 1-iii). In fact, since flows *Q* reverse periodically, they oscillate around 0. In the shear rate *τ*, oscillation periods are even doubled due to taking the absolute value of *Q* in calculating *τ*; see also Appendix A.4.

To access the long-time behavior of veins, we average out short timescales on the order of *T* ≃ 1 – 2 min corresponding to the peristaltic contractions [35]. We, thus, focus on the dynamics of the time-averaged radius 〈*a*〉 and shear rate 〈*τ*〉 on longer timescales, from 10 – 60 min (full lines in Fig. 1-iii), corresponding to growth or disassembly of the vein wall, linked to *e.g* actin fiber rearrangements [36, 37].

### B. Diverse and reproducible vein dynamics

We relate time-averaged shear rate to time-averaged vein radius and find diverse, complex, yet reproducible trajectories (Fig. 1-A/B-iv, see also Appendix 1 5 for additional datasets). To illustrate this diversity, out of 200 randomly chosen veins in the full network of Fig. 1-B, we find 80 shrinking veins, 100 stable veins, and 20 are not classifiable.

In shrinking veins, the relation between shear rate and vein adaption is particularly ambiguous. As the radius of a vein shrinks, the shear rate either monotonically decreases (pink b in Fig. 1-B-iv), or, monotonically increases (pink d), or, increases at first and decreases again (pink c). For the specimen of Fig. 1-B, out of the 80 shrinking veins, monotonic decrease is observed for 25%, monotonic increase for 40%, and non-monotonic trajectories 15% of the time. The remaining 20% of vanishing veins are unclassifiable, as their recorded trajectories are too short to allow for any classification. Out of the 12 close-up veins investigated, 4 shrink and vanish, either with monotonic or non-monotonic dynamics (see also App. 1 - figure 2).

In contrast, stable veins have a specific shear rate-radius relation: usually, stable veins perform looping trajectories in the shear rate-radius space (blue arrows in Fig. 1-A/B-iv). In the full network, these loops circle clockwise for 80% of 100 observed stable veins. Out of the 12 close-up veins investigated, 6 veins show stable clockwise feedback, 1 shows stable anticlockwise feedback, and 1 is not classifiable. Clockwise circling corresponds to an in/decrease in shear rate followed by an in/decrease in vein radius, thus, hinting at a shear rate feedback on local vein adaptation. This establishes a potential causality link between shear rate changes and vascular adaptation. In addition, the circular shape of stable vein trajectories suggests that there is a time delay between changes in shear rate and subsequent vein radius changes.

### C. Shear rate and resistance feedback alone can not account for the diversity of vein fates

We further test this potential causality link between shear rate and vein adaptation. Based on previous theoretical works [10, 27, 29–31, 33, 38, 39], we expect that the magnitude of shear rate directly determines vein fate, *i.e*. lower shear rate results in a shrinking vein. Yet, this is not corroborated by our experimental measurements. First, despite displaying comparable shear rate and vein radii at the beginning of our data acquisition, some veins are stable (blue a in Fig. 1-A/B-iv), while others vanish (pink b). We, thus, map out shear rate throughout the entire network at the beginning of our observation, see Fig. 1-B.ii. We observe that dangling ends have low shear rate, due to flow arresting at the very end of the vein (dark purple terminal veins). Yet, some dangling ends will grow (*i.e* red dot in Fig. 1-B.i.), in contradiction again with the assumption that “low shear results in a shrinking vein”. Finally, small veins located in the middle of the organism show high shear rate, yet, will vanish (yellow arrow in Fig. 1-B.ii, other examples in App. 1 - figure 4-C and App. 1 - figure 5-C). Therefore, the hypothesis that veins with low shear rate should vanish, as they cannot sustain the mechanical effort [40, 41], cannot be reconciled with our data.

Finally, also other purely geometrical vein characteristics such as vein resistance [22], 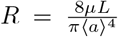, where *μ* is the fluid viscosity and *L* the vein length [42], clearly do not determine vein fate either. In fact, geometrical vein characteristics are directly related to vein radius, thus in contradiction with our observation that veins with similar radius can experience different fates (stable blue a in Fig. 1- iv and vanishing pink b). Therefore, additional feedback parameters must play a role.

## II. SHEAR RATE FEEDBACK ON INDIVIDUAL VEIN DYNAMICS OCCURS WITH A TIME DELAY

The link between shear rate feedback and vein adaptation is clearly ambiguous in our data. To understand the feedback mechanism, we now turn to modeling and indepth analysis.

### A. Vein radius adaptation in response to shear rate

Current theoretical models [6, 10, 27, 29] motivated by Murray’s phenomenological rule of minimizing dissipation [14] suggest that vascular adaptation, 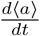, i.e. the change in time of the vein radius 〈*a*〉, is related to shear rate 〈*τ*〉 via

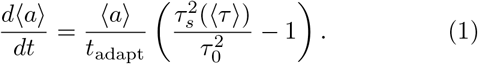

Here *τ_s_*(〈*τ*〉) is the shear rate *sensed* by a vein wall and is directly related to fluid shear rate 〈*τ*〉, in a way that we specify in the following paragraph. The parameter *t*_adapt_ is the adaptation time to grow or disassemble vein walls corresponding to fiber rearrangement [36, 37] and *τ*_0_ the vein’s reference shear rate, corresponding to a steady state regime *τ*_s_ = *τ*_0_ with constant shear rate – in agreement with Murray’s law (see Appendix 21) [14]. *t*_adapt_ and *τ*_0_ are independent variables, constants over the timescale of a vein’s adaptation, and could *a priori* vary from vein to vein, though existing models assume they do not [6, 10, 27, 29].

We here already incorporated two adaptations for our experimental system. First, we specifically indicate with 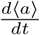 that we are interested in vascular adaptation, i.e. on long-time changes in the vein radius. In contrast, the short timescale variations 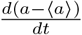 in *P. polycephalum* are driven by peristaltic contractions [35] and are not relevant for long-time adaptation. Second, we here, in contrast to all previous work, allow vein radii dynamics to potentially depend via a time delay on the shear rate, by describing radii dynamics as a function of a *sensed shear rate*, *τ_s_*(〈*τ*〉), which itself depends on the average shear rate 〈*τ*〉. We will specify this dependence in Sec. IIC.

Theoretical models differ in the precise functional dependence on shear rate on the right-hand side of Eq. 1, but agree in all using a smooth function *f* (*τ_s_*). We here employ a functional form with a quadratic scaling of the right-hand side on the shear rate 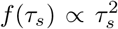 that we obtained via a bottom-up derivation from force balance on a vein wall segment in a companion work [43]. Within the force balance derivation, the cross-linked actin fiber cortex composing the vein wall responds with a force in normal direction to tangential shear and, hence, drives veins to dilate or shrink in response to shear [44, 45] (see Appendix 2 1). Experimental data measuring this anisotropic response of fibers in Refs. [45–47] suggest a quadratic dependence of the change in fibers thickness on the applied shear. This quadratic dependence is also consistent with the top-down phenomenological result of Ref. [33]. That said, our upcoming results are robust against the specific choice of *f* (*τ_s_*), as long as *f* increases with |*τ_s_*| and their exists a non-zero value of shear rate *τ*_0_ corresponding to Murray’s steady-state, *i.e*. such that *f*(*τ*_0_) = 0.

Regarding the interpretation of the sensed shear rate *τ_s_*, it is apparent from our data that the link between shear rate and radius adaptation is not immediate but occurs with a time delay. Fig. 1-iii indeed shows lag times between peaks in time-averaged shear rate and radius dynamics, ranging from 1 min to 10 min. As a result, *τ_s_* could correspond to a delayed shear rate compared to the actual one 〈*τ*〉. We turn to confirm this assumption and analyze this time delay further.

### B. Statistical analysis of the time delay between shear rate and radius dynamics

We systematically investigate the time delay between shear rate 〈*τ*〉 and vein adaptation *d*〈*a*〉/*dt*. For each vein segment, we calculate the cross-correlation between averaged shear rate 〈*τ*〉 (*t* – *t*_delay_) and vein adaptation 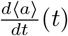 as a function of the delay *t*_delay_ (Fig. 2-A). Then, we record the value of *t*_delay_ that corresponds to a maximum (Fig. 2B). Time delays are recorded if the maximum is significant only, *i.e*. if the cross-correlation is high enough, and here we choose the threshold to be 0.5. Note, that slight changes in the threshold do not affect our results significantly. Both positive and negative time delays are recorded. Each full network data set contains more than 10000 vein segments, which allows us to obtain statistically relevant data of *t*_delay_ (Fig. 2-C and see also App. 2 - figure 2). We present additional methods to extract the time delay also in close-up networks in Appendix 2 2. Note that *t*_delay_ is different from *t*_adapt_. Although both timescales are relevant to describe adaptation in our specimen: *t*_delay_ represents the time to sense shear rate signals in vein walls; *t*_adapt_ represents the time to grow or disassemble a vein wall.

**Figure 2.**
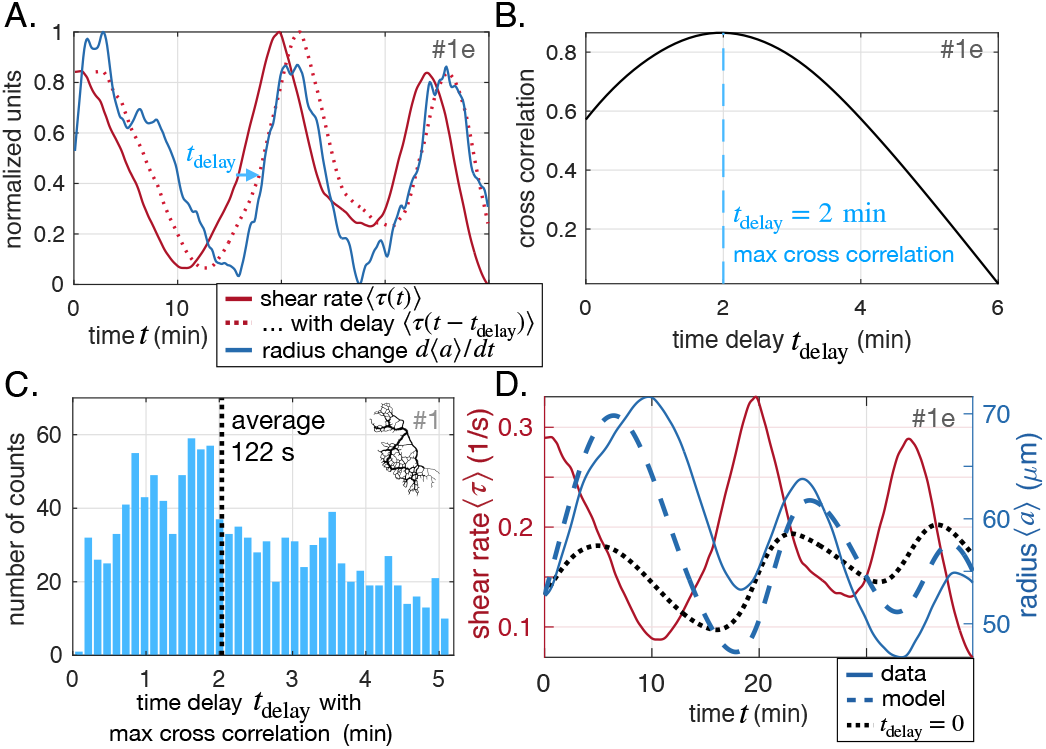
Shear rate induces vein adaptation with a time delay. (A) Principle of the cross-correlation in time between the delayed time-averaged shear rate 〈*r* (*t* – *t*_delay_)〉 and the time-averaged radius change *d*〈*a*〉(*t*)/*dt*. The plot shows the delayed curve with the best score. (B) Resulting cross-correlation for various values of *t*_delay_ and maximum extracted. (C) Statistical analysis over all veins of specimen #1 of maximal crosscorrelation with *t*_delay_, zoomed in on positive time delays, since they outnumber negative time delays by a factor of 15, see App. 2 - figure 2 for full distribution. (D) Fit result of the model Eq. (1) and Eq. (2), here, with *t*_adapt_ = 37 ± 2 min and *r*_0_ = 1.1 ±0.2 s^-1^ (heavy dashed blue). The vein investigated is the same as in (A-B) and hence we took *t*_delay_ = 2 min. The relative fitting error is *ϥ*_err_ = 0.07. The fit result of model Eq. (1) taking *r_s_* = 〈τ〉, *i.e*. with *t*_delay_ = 0 is also shown (heavy dotted black), and has error *ϥ*_err_ = 0.11.

Overall, we find 15 times more veins with positive time delays than with negative time delays for the specimen of Fig. 1-B (full time delay distribution in App. 2 - figure 2). This clearly establishes a causality link between shear rate magnitude and radius adaptation. We also find that time delays of 1 min to 3 min are quite common with an average of *t*_delay_ ≃ 2min (Fig. 2-C). We repeat the analysis over different full network specimens (App. 2 - figure 2) and close-up veins (App. 2 - figure 3) and find similar results.

While unraveling the exact biophysical origin of the time delay is beyond the scope of this work, it is important to discuss potential mechanisms. First, the typical time delay measured *t*_delay_ ~ 2 min appears close to the contraction period *T* ≃ 1 – 2 min. This is not an artifact of the analysis (see benchmark test in Appendix 2 2). Rather, it hints that the cross-linked actomyosin and contractile cortex are key players in the delay. Measured data on the contractile response of cross-linked fibers [44–46] exhibits a time delay of about 1 – 30 s for *in vitro* gels. This time delay could accumulate in much longer time delays *in vivo* [48], as is the case in our sample, and potentially reach a time delay of about 2 min. Other mechanical delays could originate from the cross-linked actomyosin gel. For example, the turnover time for actin filaments in living cells ranges from 10s to 30s [49–51], while the viscoelastic relaxation time is 100 s [52], both timescales close to our measured time delay.

### C. Model with a time delay quantitatively reproduces the data

Having clearly established the existence of a positive time delay for shear rate feedback on vein adaptation, we must radically deviate from existing models [27–31] by incorporating the measured time delay *t*_delay_ explicitly between the shear rate sensed by a vein wall *τ_s_* and fluid shear rate 〈*τ*〉. To this end, we use the phenomenological first-order equation

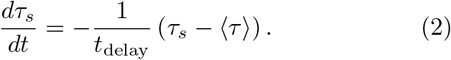

At steady state, we recover a constant shear rate 〈*τ*〉 = *τ_s_* = *τ*_0_, corresponding to Murray’s law (Appendix 2 1)[14].

We further verify that our model with the adaptation rule Eq. (1) and the time delay shear rate sensing Eq. (2) quantitatively accounts for the observed dynamics with physiologically relevant parameters. We fit our 12 closeup data sets, as well as 15 randomly chosen veins of the full network in Fig. 1-B. We take shear rate data 〈*τ*〉 (*t*) as input and fit model constants *t*_adapt_ and *τ*_0_ to reproduce radius data 〈*a*〉(*t*). Note, that *t*_adapt_ and *τ*_0_ are independent variables that vary from vein to vein, and over long timescales and between specimen [43, 53–56]. To test the robustness of model fits we employ different strategies to set the time delay *t*_delay_ before fitting. The time delay is either set to the same average value for all veins, or to the best cross-correlation value for a specific vein, or fitted with a different value for each vein, with no significant change in the resulting goodness of fit and fit parameter values.

Overall, we find a remarkable agreement between fit and data (see example in Fig. 2-D and Appendix 2 3 for more results). We find a small relative error on fitted results 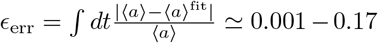. This suggests that the minimal ingredients of this model are sufficient to reproduce experimental data. Fits without the time delay yield systematically worse results, with larger fitting errors *ϵ*_err_ (see Fig. 2-D, dotted black line and Table. 2.3).

**Table I.** List of commonly used variables in our work in alphabetical order and significance. Short length scale variations correspond to variables that can vary strongly from one vein to a neighboring vein, while long length scale variations vary smoothly throughout the network. Variables have short timescale variations when they have significant variations over timescales much smaller than the peristaltic contractions *T* ≃ 1 — 2 min; and long timescale variations if they vary over longer timescales corresponding to vascular adaptation and rearrangement.

| Variable | Significance | Length scale variations | Timescale variations |
| --- | --- | --- | --- |
| $a$ | radius of a vein | short | short and long |
| $\epsilon$ | relative contraction amplitude | – | – |
| $L$ | length of a vein | – | – |
| $\mu$ | inner fluid viscosity | long | long |
| $\omega = 2\pi/T$ | contraction frequency | – | – |
| $P$ | inner fluid pressure | long | short and long |
| $P_0$ | atmospheric pressure | – | – |
| $Q$ | fluid flow pervading a vein | short | short and long |
| $Q_{\text{in}}$ | fluid flow generated by peristaltic contractions in a vein | short | short and long |
| $Q_{\text{net}}$ | fluid flow generated by the rest of the network | short | short and long |
| $R$ | resistance of a vein, $R = 8\mu L/\pi a^4$ | short | short and long |
| $R_{\text{net}}$ | resistance of the rest of the network attached to a vein at both vein ends | short | short and long |
| $\tau$ | shear rate on a vein's inner wall | short | short and long |
| $\tau_s$ | sensed shear rate for adaptation | short | long |
| $\tau_0$ | steady state shear rate, $\tau_0 \sim \tau_{\text{active}} - \langle P - P_0 \rangle/\mu$ | long | long |
| $\tau_{\text{active}}$ | Active contribution to $\tau_0$, due to energetic consumption from the actomyosin cortex to maintain contractions | long | long |
| $t$ | time | – | – |
| $t_{\text{adapt}} \simeq 10 - 100 \text{ min}$ | long-time adaptation timescale | long | long |
| $t_{\text{delay}} \simeq 2 \text{ min}$ | delay between adaptation and shear rate | long | long |
| $T \simeq 1 - 2 \text{ min}$ | peristaltic contraction period | – | – |
| $\langle X \rangle$ | long-time average of $X$ | – | long |

In all samples, fitting parameters resulted in physically reasonable values. We found *t*_adapt_ ≃ 10 – 100 min corresponding to long timescale adaptation of vein radii. Note again, the physical difference between the time to adapt vein radius *t*_adapt_ and the time delay to sense shear rate *t*_delay_ also translates to orders of magnitude differences with *t*_delay_ ≃ 2 min and *t*_adapt_ ≃ 10 – 100 min. This 10 – 100 min is indeed the timescale over which we observe significant adaptation. Reorganization of biological matter occurs on similar timescales in other comparable systems, from 15 min for individual cells to several days for blood vasculature [50, 57].

When examining fit results of the target shear rate *τ*_0_ it is *a priori* hard to estimate which values to expect since *τ*_0_ is only reached at steady state. Yet, in our continuously evolving specimen, we never reach steady state and, hence, can not measure *τ*_0_. However, we can compare *τ*_0_ to shear rate values measured in our specimen and find that they are consistently of the same order of magnitude. Finally, we find that our model yields better results if we fit the data over intermediate time frames (15 min to 40 min), exceeding results of fitting over longer time frames (40 min to 100 min). This is in line with our theoretical expectation [43] that *t*_adapt_ and *τ*_0_ change over long timescales, since they depend on physical parameters that also change over long timescales, in particular in response to network architecture changes. Since veins typically vanish over 15 min to 40 min and, hence, significant network changes occur over exactly that timescale, *t*_adapt_ and *τ*_0_ are no longer constant for time frames ≳ 40 min.

While we have focused so far on timescales of individual vein adaptation, we now aim to understand how their individual disparate fates arise. We will show that the origin of different fates resides in the evolution of the rest of the network.

## III. RELATIVE RESISTANCE AND PRESSURE DETERMINE VEIN FATE WITHIN A NETWORK

### A. Stable and unstable vein dynamics are predicted within the same model

To capture the impact of the entire network on the dynamics of a single vein modeled by Eqs. (1–2), we must specify the flow-driven shear rate 〈*τ*〉. Since 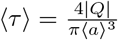, it is sufficient to specify the flow rate *Q* in a vein. *Q* is coupled to the flows throughout the network by conservation of fluid volume through Kirchhoff’s laws, and is, therefore, an indirect measure of network architecture.

We, here, consider the most common vein topology of a vein connected at both ends to the remaining network, more specialized topologies follow in Sec. III C. The network is then represented by a vein of equivalent resistance *R*_net_ parallel to the single vein of 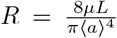 considered within an equivalent flow circuit, see Fig. 3-A. *R*_net_ is the equivalent resistance corresponding to all the resistances making up the rest of the network, obtained with Kirchhoff’s laws (see examples in Appendix 31). *R*_net_ is therefore integrating the network’s architecture. Such a reduction of a flow network to a simple equivalent flow circuit is always possible due to Norton’s theorem [58].

**Figure 3.**
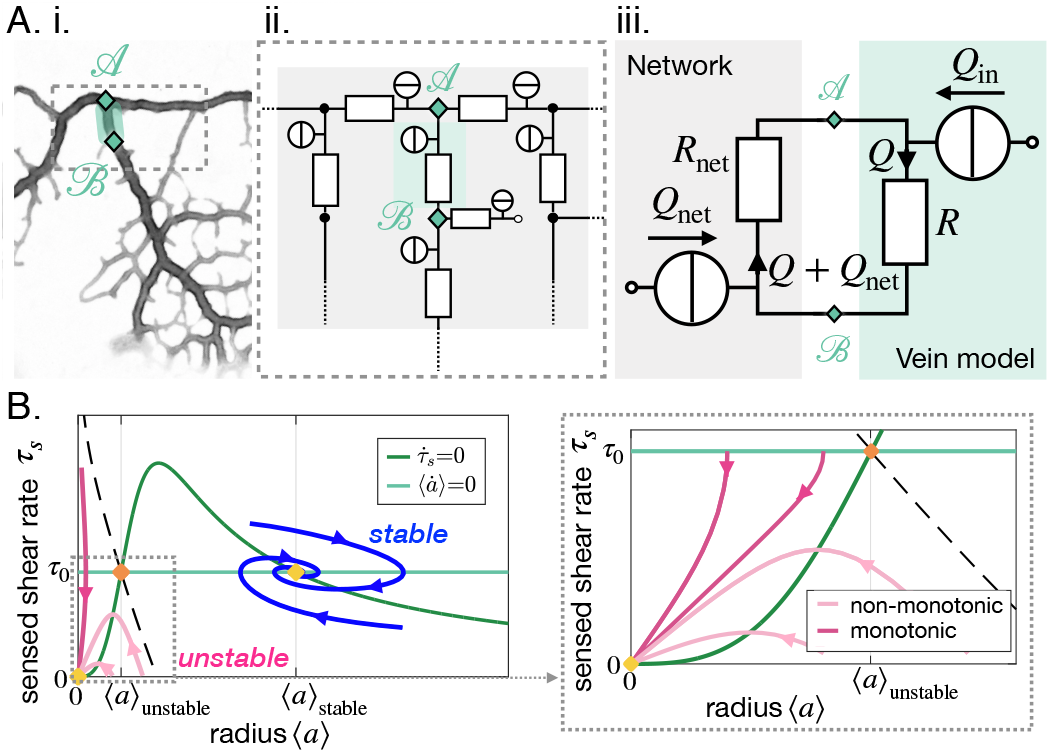
Stable and unstable vein dynamics are predicted within the same model. (A) Translation of (i) a bright field image of specimen into (ii) vein networks; each vein is modeled as a flow circuit link. (iii) Reduction of (ii) via Northon’s theorem into an equivalent and simplified vein flow circuit consisting of a flow source *Q*_in_ (due to vein’s pumping) and a resistor *R* (viscous friction). The rest of the network is modeled by an equivalent circuit with flow source Qnet = – *Q*_in_ and resistor *R*_net_. *Q* flows through the vein. (B) (Left) Time-averaged sensed shear rate *τ_s_* versus radius from (1)-(3) with fixed points and typical trajectories. The green lines correspond to stationary solutions for *τ_s_* or 〈*a*〉. The blue lines correspond to stable trajectories and the pink lines to unstable ones. (Right) Zoom of the phase space corresponding to shrinking veins, including monotonic and non-monotonic trajectories.

The time-averaged net flow generated by the vein contractions is 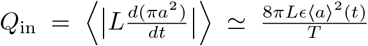 where *ϵ* is the relative contraction amplitude. The absolute values in this definition are used to measure the *net* flow. *Q*_in_ thus measures the mass exchanges between the network and the vein. As mass is conserved, this results in an inflow of *Q*_net_ = – *Q*_in_, into the rest of the network. Within the vein, a total flow rate *Q* circulates – see Fig. 3-A. ii. The flow rate *Q* through the vein follows from Kirchhoff’s second law: *QR* = – (*Q* + *Q*_net_)*R*_net_. We, thus, obtain that the time-averaged shear rate in the vein is

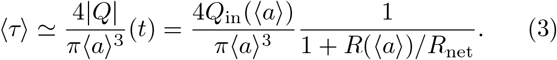

The coupled dynamics of {〈*τ*〉, *τ_s_*, 〈*a*〉} are now fully specified through Eq. (1–3). To simplify our analysis, we now explore the reduced system {*τ_s_*, 〈*a*〉} by replacing 〈*τ*〉 in Eq. (2) by its expression in Eq. (3). Using standard tools of dynamical systems theory, see Appendix 3 3, we now characterize the typical trajectories predicted within the model.

Our dynamic system {*τ_s_*, 〈*a*〉} reproduces the key features of the trajectories observed experimentally. We find two stable fixed points at (0, 0) and (*τ*_0_, 〈*a*〉_stable_(*R/R*_net_, *τ*_0_)), and one unstable fixed point at (*τ*_0_, 〈*a*〉_unstable_(*R*/*R*_net_, *τ*_0_)) (see Fig. 3-B). The stable fixed point with finite radius, (*τ*_0_, 〈*a*〉_stable_) corresponds to Murray’s steady state. The set of fixed points was also found in a related theoretical study that investigates a phenomenological model resembling Eq. (1), yet without any time delay, and examining the stability of a vein connected to a pressure source and a another resistance [27]. This suggests that the presence of the three fixed points is universal. Furthermore, we find similar dynamical trajectories in the {*τ_s_*, 〈*a*〉} as those observed experimentally. Trajectories spiral in the clockwise direction near the stable fixed point (*τ*_0_), 〈*a*〉_stable_) (blue in Fig. 3-B) and veins shrink with monotonic (dark pink in Fig. 3-B) or with non-monotonic shear rate decrease (light pink Fig. 3-B). The dynamics of 〈*τ*〉 are then closely related to that of *τ_s_*.

### B. Relative resistance and pressure control vein fate

Analysis of the vein network model as a dynamic system, Eqs. (1–3), clearly highlights that different vein fates may occur depending on the value of the relative resistance *R*/*R*_net_ and on the value of the target shear rate *τ*_0_ for that specific vein. We will, therefore, now investigate their values throughout the network more carefully.

Before proceeding, we must specify the meaning of the target shear rate *τ*_0_. The force balance derivation in Ref. [43] finds that the shear rate reference *τ*_0_ is related to the local fluid pressure *P*, as *τ*_0_ ~ *τ*_active_ – 〈*P* – *P*_0_〉/*μ* (see short derivation in Appendix 21). Here *P* – *P*_0_ characterizes the pressure imbalance between the fluid pressure inside the vein, *P*, and the pressure outside, *P*_0_, namely the atmospheric pressure. We recall that *μ* is the fluid viscosity. Finally, *τ*_active_ = *σ*_active_/*μ* is a shear rate related to the active stress *σ*_active_ generated by the actomyosin cortex [59, 60]. The active stress sustains the contractile activity of the vein, and is, therefore, an indirect measure of the metabolic or energetic consumption in the vein. The local pressure P results from solving Kirchhoff’s law throughout the network. It is, therefore, indirectly integrating the entire network’s morphology. Hence, not only *R/R*_net_ but also *τ*_0_ is a flow-based parameter, integrating network architecture.

In our experimental full network samples, we can calculate both the relative resistance *R*/*R*_net_ and the local pressure *P*, and its short time-averaged counterpart 〈*P*〉, up to an additive constant (see Fig. 4). We find that pressure maps of 〈*P*〉 are mostly uniform, except towards dangling ends where relevant differences are observed (Fig. 4-A). Hence, particularly in dangling ends, veins with similar shear rate *τ* may suffer different fates, as described through Eq. (1). This is a radical shift compared to previous theoretical works which consider that *τ*_0_ is a constant throughout the network [10, 27, 29–31, 33, 38, 39].

**Figure 4.**
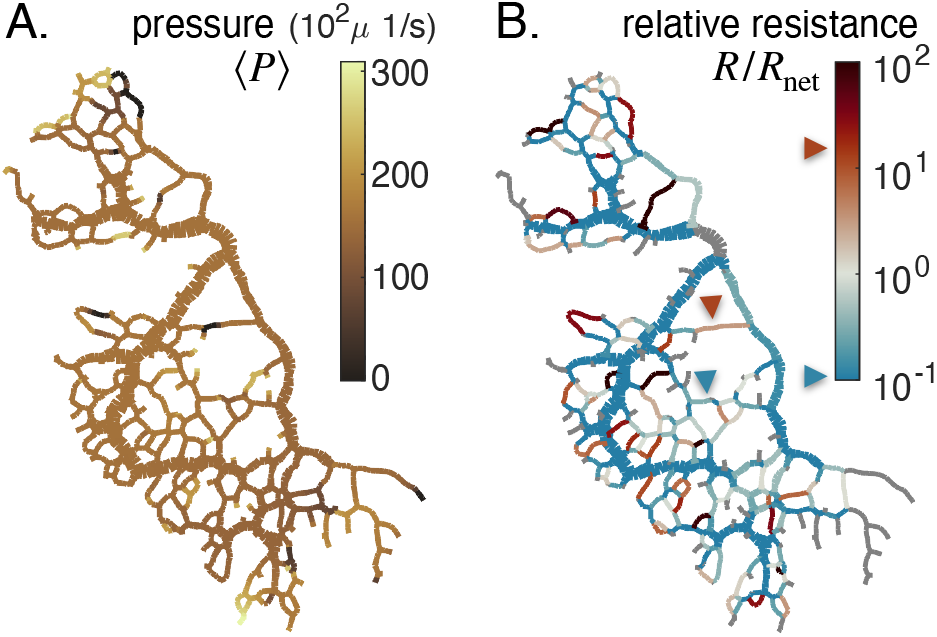
Feedback parameters integrate the network’s architecture and provide information on vein relative location. Full network maps of the same specimen as in Fig. 1B, at the beginning of the observation, of (A) the average fluid pressure in a vein 〈*P*〉 and (B) of the relative resistance *R/R*_net_. The fluid pressure 〈*P*〉 is defined up to an additive constant. Grey veins in (B) correspond to bottleneck veins or dangling ends for which *R*_net_ can not be defined. The color scales indicate the magnitude of each variable in each colored vein. For example, in (B), the red arrow indicates a vein with large relative resistance *R/R*_net_.

The relative resistance *R/R*_net_ varies over orders of magnitude (Fig. 4-B), with values that are not correlated with vein size (see Appendix 1 - figure 6). Rather, *R/R*_net_ indicates how a vein is localized within the network compared to large veins that have lower flow resistance and that serve as highways for transport. For example, a small vein immediately connected to a highway will show a large value of *R/R*_net_. In this case among all possible flow paths that connect the vein’s endpoints, there exists a flow path that consists only of highways, and therefore we expect *R* ≫ *R*_net_ (see red arrow in Fig. 4-B). In contrast, a similarly small vein yet localized in between other small veins, further away from highways, will show a smaller value of *R*/*R*_net_. In this latter case, all flow paths have to pass through small nearby veins and, hence, have high resistance *R*_net_ 〈 *R* (see blue arrow in Fig. 4-B). *R/R*_net_, therefore, reflects the relative cost to transport fluid through an individual vein rather than through the rest of the network.

The relative resistance *R/R*_net_ is, thus, a natural candi-date to account for individual vein adaptation: it measures the energy dissipated by flowing fluid through an individual vein, *Q*^2^*R*/2, compared to rerouting this flow through the rest of the network, *Q*^2^*R*_net_/2. Hence, we may expect that when in a given vein *R* > *R*_net_, it is energetically more favorable to flow fluid through the rest of the network and hence to shrink the vein. Reciprocally, if *R* < *R*_net_, we expect that the vein is stable. Analyzing our equations gives further support to this intuitive rule. When *R* ≫ *R*_net_, from Eq. (3), we may expect 〈*τ*〉 to be relatively small, in particular, small relative to the vein’s specific steady state *τ*_0_ and hence via Eq. (1) the vein would likely shrink. Reciprocally, if *R* ≪ *R*_net_, we may expect 〈*τ*〉 to be relatively large compared to its specific *τ*_0_, and hence the vein is stable. Yet, since *R/R*_net_ is nondimensional, it can provide more systematic insight than 〈*τ*〉, since *τ*_0_ is not known *a priori*. Notice that the red arrow in Fig. 4-B presents a shrinking vein that indeed verifies *R* > *R*_net_. However, according to shear rate measures (see yellow arrow in Fig. 1-B.ii), the shear rate is large in that vein, preconditioning the vein to grow, according to previous works [10, 27, 29–31, 33, 38]. We can therefore show why occasionally, veins at high shear rate shrink, and veins at low shear rate grow by highlighting that *R/R*_net_, beyond shear rate, is crucial to predict vein fate.

Our aim is now to investigate, in more detail, how these novel feedback parameters integrating network architecture, the relative resistance *R/R*_net_ and the local pressure P via the target shear rate, control vein dynamics on the basis of three key network topologies of a vein.

### C. Specific vein fates: dangling ends, parallel veins and loops

#### Dangling ends are unstable: disappearing or growing

As observed in our data, dangling ends are typical examples of veins that can start with very similar shear rate and radius and yet suffer radically different fates (Fig. 1-B.i and ii, Fig. 5-A). Dangling ends either vanish or grow but never show stably oscillating trajectories.

**Figure 5.**
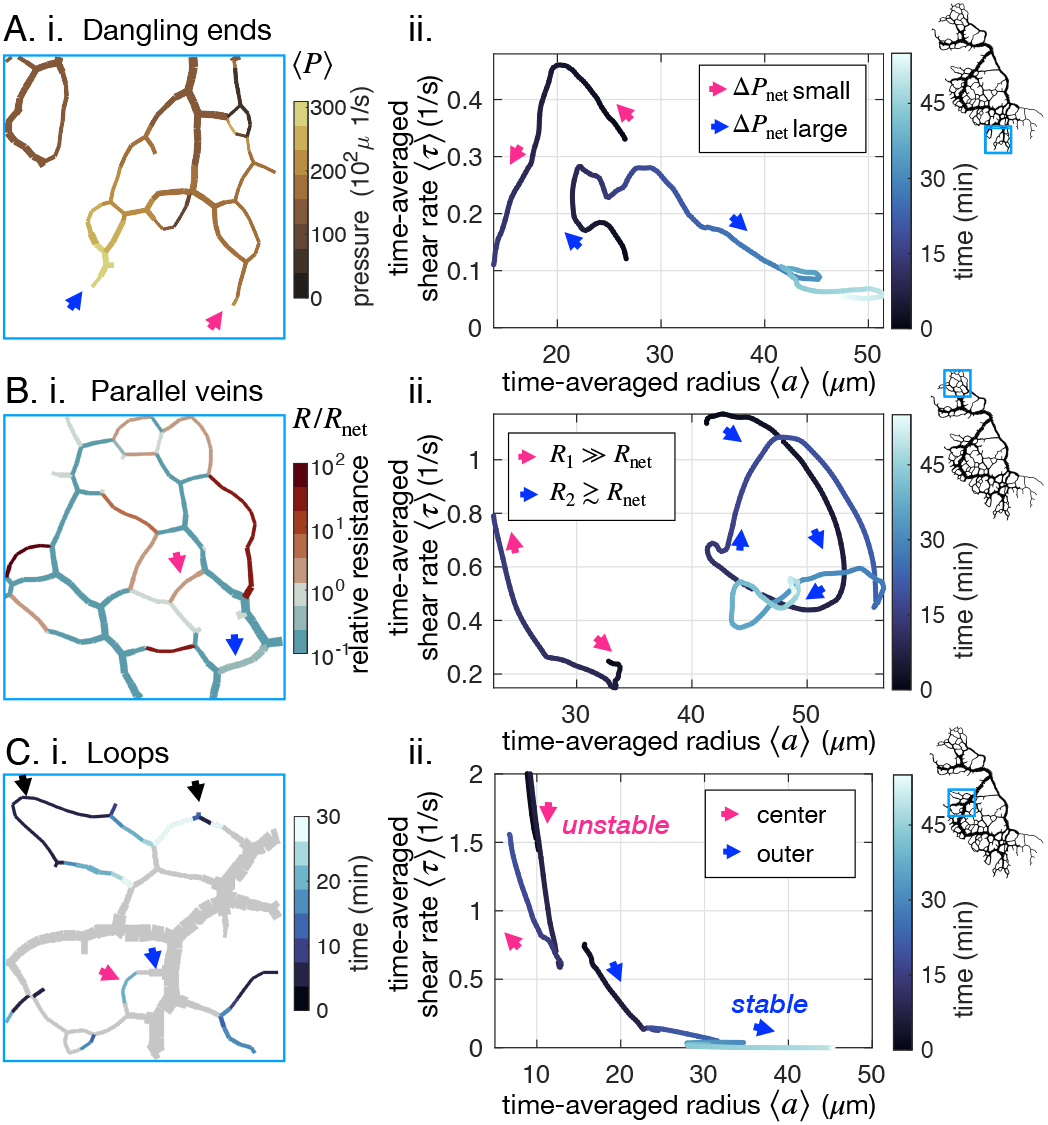
Network architecture controls vein fate as exemplified in three cases. (A-C) (i) determining factors mapped out from experimental data for the specimen of Fig. 1B and (ii) typical trajectories from data. All pink (respectively blue) arrows indicate shrinking (respectively stable or growing) veins. (A) (ii) Dangling ends either vanish or grow indefinitely, coherently with (i) the relative local pressure 〈*P*〉. Arrows point to veins initially similar in size (~ 23 μm). (B) Parallel veins are unstable: one vanishes in favor of the other one remaining (ii), coherently with (i) its relative resistance, *R/R*_net_, being higher. (C) Loops first shrink in the center of the loop (ii) – *i.e*. from the point furthest away from the nodes connecting it to the rest, of the network – as evidenced by focusing on (i) the time of vein segment vanishing. Black arrows point to other loops also vanishing from the center. For all graphs on the left, the color scales indicate the magnitude of each variable in each colored vein.

Topologically, and unlike the middle vein considered in Fig. 3-A, dangling ends are only connected to the rest of the network by a single node. Therefore, the relative resistance Rnet cannot be calculated in a dangling end and cannot play a role. The shear rate in a dangling end is simply 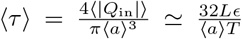. Using this expression instead of Eq. (3) and analyzing the dynamical system with Eq. (1–2), we find that veins can only shrink or grow (see Appendix 41). Furthermore, *τ*_0_ determines the threshold for growth over shrinkage. Since *τ*_0_ ~ *τ*_active_ – 〈*P* – *P*_0_〉/*μ* a large 〈*P*〉 decreases *τ*_0_. Hence, the model predicts that a larger pressure at a dangling end facilitates growth.

We observe for the example of Fig. 5-A that large values of 〈*P*〉 indeed appear to favor growth, and small values prompt veins to vanish. This agrees with physical intuition: when a vein is connected to a large input pressure, one expects the vein to open up. Notice, however, that here the mechanism is subtle. The shear rate itself is not large. Rather, the shear rate threshold to grow is lowered by the high local pressure. Local pressure is thus connected to dangling end fate: it is a prime example of the importance of *integrating network architecture*.

#### Competition between parallel veins decided by relative resistance

Parallel veins are another example in which initially very similar and spatially close veins may suffer opposite fates; see Fig. 5-B. Often, both parallel veins will eventually vanish, yet what determines which vanishes first?

To investigate this situation we can simply extend the circuit model of Fig. 3-A with another parallel resistance, corresponding to the parallel vein (Appendix 4 2). We then have two veins with respective resistance say *R*_1_ and *R*_2_. We can analyze the stability of this circuit with similar tools as in Sec. III A. We find that if one vein’s relative resistance is larger than the other one’s, say for example *R*_1_/*R*_net,1_ > *R*_2_/*R*_net,2_, then vein 1 vanishes in favor of the other vein 2 as previously predicted in simpler scenarios for steady states [27]. Exploring *R/R*_net_ in our full network (Fig. 5-B), we find that a vein with a large relative resistance *R/R*_net_ > 1 will vanish. In contrast, a nearby, nearly parallel vein with *R/R*_net_ ≃ 1 will remain stable.

The relative resistance *R/R*_net_ is thus a robust predictor for locally competing veins. Although it is connected to shear rate, as highlighted through Eq. (3), there are clear advantages to the investigation of *R/R*_net_ over the shear rate itself: *R/R*_net_ is straightforward to compute from global network architecture as it does not require to resolve flows, and it is non-dimensional.

#### Loops shrink first in the middle

Finally, loopy structures *i.e*. a long vein connected at both ends to the remaining network, are often observed in *P. polycephalum*. Surprisingly, we experimentally observe loops to start shrinking in their very middle (Fig. 5-C, App. 1 - figure 4-F and App. 1 - figure 5-F) despite the almost homogeneous vein diameter and shear rate along the entire loop. This is all the more surprising as quantities such as 〈*P*〉 and *R/R*_net_ are also similar along the loop.

This phenomenon again resides in the network architecture, and we can rationalize it with an equivalent flow circuit (see Appendix 4 3). When a vein segment in the loop shrinks, mass has to be redistributed to the rest of the network. This increases shear rate in the outer segments, preventing the disappearance of the outer segments of the loop. Once the center segment has disappeared, both outer segments follow the dynamics of dangling ends. Their fate is again determined by network architecture, through the local pressure 〈*P*〉 in particular.

Importantly, we find that as soon as a vein disappears, the network’s architecture changes: flows must redistribute, and vein connections are updated. Hence, an initially stable vein may become unstable. Vein fates, thus, dramatically evolve over time.

## IV. SINGLE VANISHING VEIN TRIGGERS AN AVALANCHE OF VANISHING EVENTS AMONG NEIGHBORING VEINS

After focusing on individual vein dynamics, we now address global network reorganization. Observing a disappearing network region over time, we find that veins vanish sequentially in time (Fig. 6-A and B). Inspired by the importance of relative resistance for parallel veins, we here map out relative resistance *R/R*_net_ at subsequent time points in an entire region (Fig. 6-A). At the initial stage (Fig. 6-A, 2 min), the majority of veins are predicted to be stable with a relative resistance *R/R*_net_ < 1. As expected, the few veins with high relative resistance (red arrows in Fig. 6-A, 2 min) indeed vanish first (black crosses in Fig. 6-A, 5 min).

**Figure 6.**
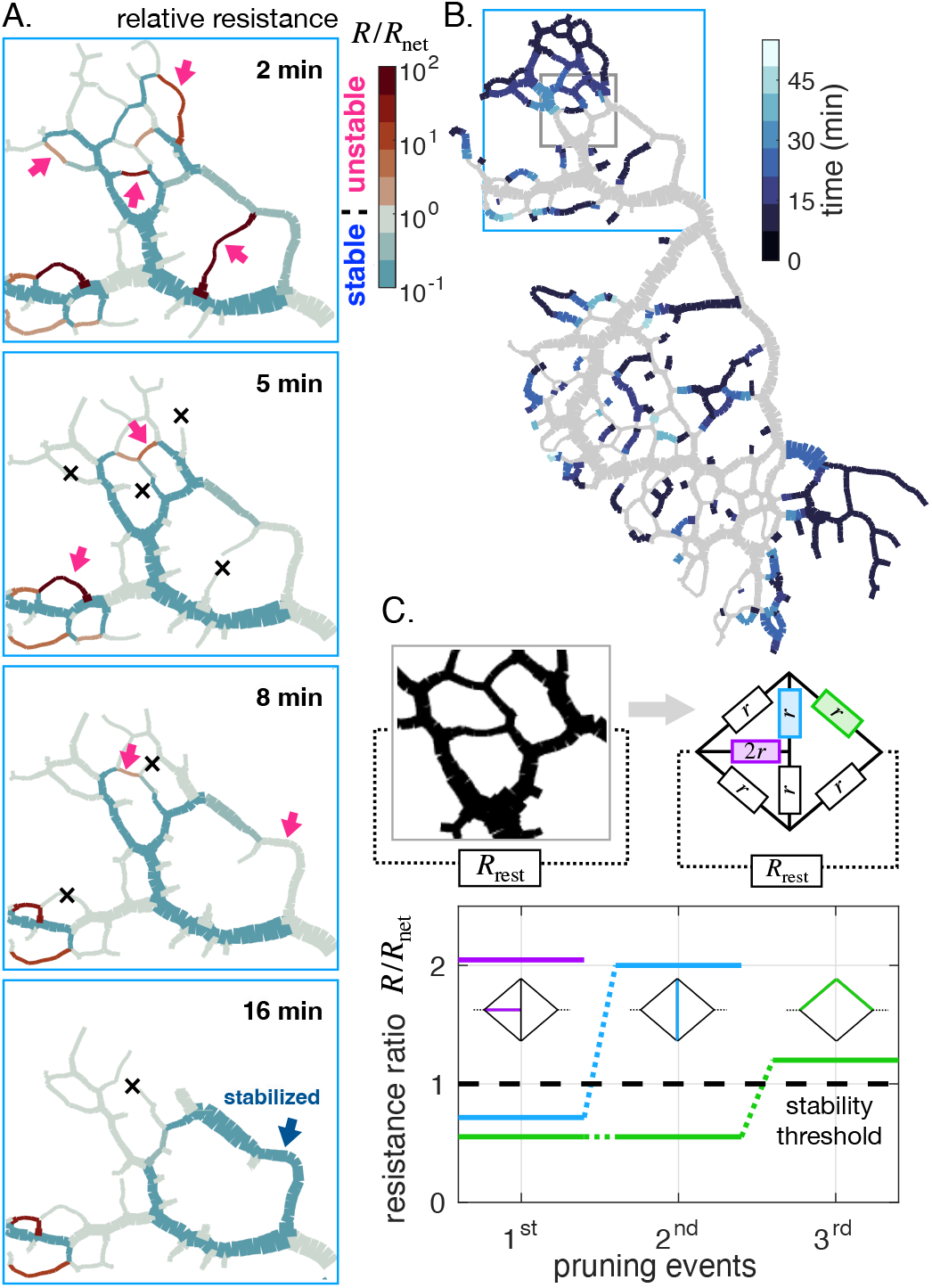
Avalanche of sequentially vanishing veins. (A) Time series of network reorganization. Each vein is colored according to the ratio between the resistance of an individual vein R and the rest of the network *R*_net_ in each vein. Red arrows highlight vanishing veins in the experiment; black crosses indicate veins that disappeared within the previous time frame. Veins for which the relative resistance cannot be calculated, such as dangling ends, are plotted with *R/R*_net_ = 1. (B) Map indicating vanishing vein events, with veins colored according to their disappearance time reported in the color scale. Gray veins will remain throughout the experiment. (C) Dynamics of the relative resistance of the three color-coded veins within a minimal network, inspired by the highlighted gray region of the network in (B). Vein resistances are chosen as *R* = *r* except for a perturbed vein for which *R* = 2r. *R*_rest_ represents the rest of the network relative to the region.In this model, a vein vanishes if its individual relative resistance *R/R*_net_ > 1. The disappearance of veins sequentially increases the relative resistance of neighboring veins, making them unstable. Here *r/R*_rest_ = 0.1, yet similar behavior was obtained consistently over a wide range of *r* values.

As a consequence of veins vanishing, the local architecture is altered, and the relative resistance, through *R*_net_, changes drastically. Veins that were stable before are now predicted to be unstable. This avalanche-like pattern, in which individual vanishing veins cause neighboring veins to become unstable, repeats itself until the entire region disappears in less than 15 min (Appendix 1 - figure 4-F and App. 1 - figure 5-F show similar avalanches in other specimens). Note that a vanishing vein may rarely also stabilize a previously unstable vein (Fig. 6-A, 16 min, blue arrow).

The fundamental origin of these avalanches of vanishing veins can be narrowed down again to network architecture. We explore a model network region, inspired by a region in an actual specimen (Fig. 6-C). We simplify the investigation by considering the region is made of a few veins of similar resistance *r* connected to the rest of a network, represented by an overall equivalent resistance *R*_rest_. *R*_rest_ represents the rest of the network relative to the region, distinct from *R*_net_, which is relative to a single vein. We precondition all veins to be stable, assuming that for each vein its relative resistance *R/R*_net_ ≾ 1. Since in our model network for each vein we approximately have *R/R*_net_ ~ *r/R*_rest_ this prescribes the initial values of *r/R*_rest_ ≾ 1.

We now perturb a vein slightly, *e.g*. with a smaller radius, and therefore with a slightly higher resistance, say 2r (purple in Fig. 6-C). The perturbed vein’s relative resistance thus may become greater than 1, making the vein unstable. As the vein vanishes, two network nodes are removed, and individual veins previously connected through the node now become a single *longer* vein. A longer vein has a higher hydraulic resistance. Hence, the “new” longer vein also becomes unstable (blue in Fig. 6-C). Once it vanishes, in turn, another neighboring vein becomes longer and unstable (green in Fig. 6-C). Reciprocally, vein growth and parallel vein disappearance can – more rarely – decrease *R/R*_net_, and in turn, stabilize a growing vein, as highlighted by the blue arrow in Fig. 6-A at 16 min.

In our simple mechanistic model, the series of events follows an avalanche principle similar to that observed in our experiments: a vanishing vein disturbs local architecture. This modifies the relative resistance of nearby veins and hence their stability. The avalanche of disappearing veins eventually results in the removal of entire network regions.

## DISCUSSION

We here report highly resolved data of spontaneous network reorganization in *P. polycephalum* in which both individual vein dynamics and fluid flows pervading veins are quantified simultaneously. We observe disparate vein dynamics originating from shear-driven feedback on vein size. Strikingly, shear-driven feedback occurs with a time delay ranging from 1 min to 3 min. Our vein network model challenges previous concepts showing that vein fate is not only determined through shear rate magnitude but also through parameters that integrate network architecture via fluid flow. In particular, dangling end fate is connected to local fluid pressure 〈*P*〉, with larger pressures stabilizing dangling ends. Inner network vein fate is tightly determined by the vein’s resistance relative to the resistance to fluid flow through the rest of the network, *R/R*_net_. When *R/R*_net_ > 1 (reciprocally *R/R*_net_ < 1), this preconditions the vein to shrink (respectively to grow or be stable). While *R/R*_net_ is directly related to shear, it can be easily computed from network morphology, *without* needing to resolve flows. Both relative resistance *R/R*_net_ and local pressure 〈*P*〉 are based on fluid flow physics and are indirect measures of the entire network architecture. Yet, network architecture strongly depends on time. As unstable veins vanish, the relative architecture of changes, inducing avalanches of vanishing veins, resulting in significant spontaneous reorganization.

While our experimental investigation is specific to *P. polycephalum*, we expect that the two key concepts unraveled here, time delay and network architecture governing vein fate through relative resistance and fluid pressure, may very well be at play in other vascular networks. First, the ubiquity of delayed shear rate feedback, beyond the contractile response of actomyosin, suggests that a diversity of vein dynamics (circling, non-monotonic) may also occur in other vascular networks. In fact, also the turnover time for actin filaments in living cells ranges from 10 s to 30 s, close to our measured time delay [49–51]. Other pathways, such as chemical pathways for sheared endothelial cells in blood vasculature, are processed with a time delay of a few minutes [61–63], while reorganization occurs on longer timescales ranging from 15 min for individual cells to several days for blood vasculature [50, 57].

Second, network architecture feedback, through relative resistance and pressure, is connected to the laminar flows pervading the network. Thus, our perspective could be extended to other networks where laminar flows are an essential building block, in essence, to the diversity of networks where Murray’s law holds at steady state [17–21]. Particularly, our insight suggests simple parameters to map out, such as the purely geometrical relative resistance. Likely these parameters, which integrate network architecture, may explain discrepancies between shear rate and network reorganization in other vascular networks [3, 22–25].

Notably, imaging of biological flow network as a whole is, as of now, a rare feature of our experimental system that enabled us to unravel the importance of the network architecture for vein fate. Yet, we are hopeful that future theoretical work may allow for vein fate prediction with relative resistances determined only with partial information of a network’s architecture, with sufficient accuracy. At the same time, novel experimental techniques now open up the way for *in toto* imaging of vascular systems and quantitative assessment of dynamics [64].

The fact that pervading flows and network architecture are so intermingled originates in the simple physical principle that flows are governed by Kirchhoff’s laws at nodes, and hence “autonomously” sense the entirety of the network’s architecture. Yet, Kirchhoff’s laws are not limited to flow networks, but also govern electrical [65], mechanical [66–69], thermal [70] and resistor-based neural networks [71, 72]. Having the physics of Kirchhoff-driven self-organization at hand may thus pave the way for autonomous artificial designs with specific material [66, 67] or learning properties [65, 71, 72].

## ACKNOWLEDGMENTS

The authors are indebted to Charles Puelz, Emilie Verneuil, and Agnese Codutti for enlightening discussions. S.M. was supported in part by the MRSEC Program of the National Science Foundation under Award Number DMR-1420073. This work was supported by the Max Planck Society and has received funding from the European Research Council (ERC) under the European Union’s Horizon 2020 research and innovation programme (grant agreement No. 947630, FlowMem).

## Appendix 1: Preparation, imaging, and general data analysis

Microscopic images of all the specimens used for this study are available as movies in MP4 format. Numerical data analysis available upon request to the corresponding author.

### 1. Preparation and Imaging of *P. polycephalum*

*P. polycephalum* (Carolina Biological Supplies) networks were prepared from microplasmodia cultured in liquid suspension in culture medium [73, 74]. For the full network experimental setup, as in Fig. 1-B of the main text, microplasmodia were pipetted onto a 1.5% (w/v) nutrient free agar plate. A network developed overnight in the absence of light. The fully grown network was trimmed in order to obtain a well-quantifiable network. The entire network was observed after 1 h with a Zeiss Axio Zoom V.16 microscope and a 1x/0.25 objective, connected to a Hamamatsu ORCA-Flash 4.0 camera. The organism was imaged for about an hour with a frame rate of 10 fpm.

In the close-up setup, as in Fig. 1-A of the main text, the microplasmodia were placed onto a 1.5% agar plate and covered with an additional 1 mm thick layer of agar. Consequently, the network developed between the two agar layers to a macroscopic network which was then imaged using the same microscope setup as before with a 2.3x/0.57 objective and higher magnification. The high magnification allowed us to observe the flow inside the veins for about one hour. Typical flow velocities range up to 1 mms^-1^ [75]. The flow velocity changes on much longer timescales of 50s to 60s. To resolve flow velocity over time efficiently 5 frames at a high rate (typically 60 ms, detailed frame rates are specified for each Video) were imaged separated by a long exposure frame of about 2 s. As different objectives were required for the two setups, they could not be combined for simultaneous observation. Typically the longer exposure frame appears as a bright flash in the Videos. The 12 close-up data sets are indexed #*A* – *L* consistently in the main text and Appendix.

### 2. Image Analysis

For both experimental setups, image analysis was performed using a custom-developed MATLAB (The MathWorks) code. This procedure extracts the entire network information of the observed organism [73]: single images were binarized to identify the network’s structure, using pixel intensity as well as pixel variance information, extracted from an interval of images around the processed image. As the cytoplasm inside the organism moves over time, the variance gives accurate information on which parts of the image belong to the living organism and which parts are biological remnants. The two features were combined and binarized using a threshold. The binarized images were skeletonized and the vein radius and the corresponding intensity of transmitted light were measured along the skeleton. The two quantities were correlated according to Beer-Lambert’s law and the intensity values were further used as a measure for vein radius, as intensity provides higher resolution. For the imaging with high magnification, in addition to the network information, the flow field was measured using a particle image velocimetry (PIV) algorithm inspired by [76–78], see Fig. 1-A.ii of the main paper. The particles necessary for the velocity measurements are naturally contained within the cytoplasm of *P. polycephalum*.

### 3. Flow calculation from vein contractions

Building on the previous image analysis, we used a custom-developed MATLAB (The MathWorks) code to calculate flows within veins for the full networks, based on conservation of mass. The algorithm follows a two stage process.

First, the network structure obtained from the images was analyzed to construct *a dynamic* network structure. This structure consists in discrete segments that are connected to each other at node points. At every time point, the structure can evolve according to the detected vein radii: if a radius is lower than a certain threshold value, the corresponding segment vanishes from the structure. Segments which are isolated due to vanishing segments are also removed. We carefully checked by eye that the threshold levels determining when a segment vanished agreed with bright-field observations. Note that we do not account for entirely new segments in the dynamic structure. As no substantial growth occurs in our data, this is a good approximation.

Second, flows and pressure in each segment were calculated building on Ref. [8]. We formalize this step briefly. Let *n* and *p* be two indices to describe node *n* and node *p* connected by a segment say *i*. In each segment, there is an unknown inflow from neighboring segments *Q*_0,*np*_. There is also added flow arising due to periodic contractions 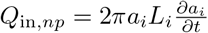 where *α_i_* denotes the radius of segment *i* and *L_i_* is the length of the vein. Note that all flows are given directed from node *n* to node *p*. As a result the flow arriving from segment *i* at node *p* is simply *Q*_0,*np*_ + *Q*_in,*np*_. According to Kirchhoff laws, at each node in the network, at each time point, the total incoming flux from each segment has to be zero

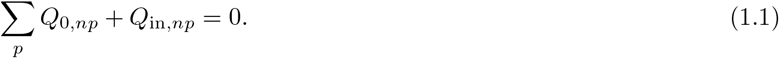

This can be rewritten

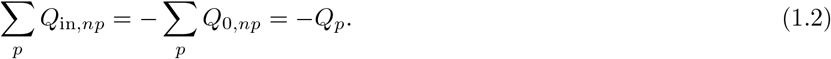

where *Q_p_* are the new unknowns. Since Poiseuille law holds, the *Q*_0,*np*_ are given by

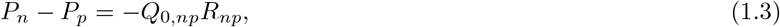

where *P_n_* is the local pressure at node *n* (respectively *P_p_* at node *p*) and *R_np_* is the hydraulic resistance of a vein such that 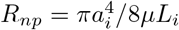. Hence

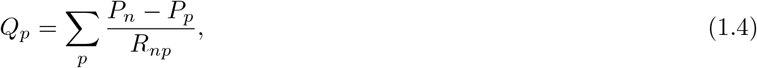

which is a linear equation of the form 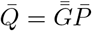 where 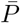 is the vector of pressure at each node in the network, similarly 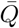 is the vector of unknown inflows at each node, and 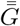 is a matrix of inverse resistances taking into account the architecture of the network. We can invert this equation to obtain the values of pressure at network nodes. Then we calculate the inflow from neighboring veins through Eq. (1.3). Finally, we obtain pressure in segment *i* as *P_i_* = (*P_n_* + *P_p_*)/2.

Compared to Ref. [8], we introduced two major additions. On the one hand, the actual live contractions *a_i_*(*t*) are used, as detected from sequential images. To ensure that Kirchhoff’s laws are solved with a good numerical accuracy, the radius traces *a_i_*(*t*) were (1) adjusted at each time so that overall cytoplasmic mass is conserved (mass calculated from image analysis varied by less than 10% over the analysis time) and (2) overdiscretized in time by adding 2 linearly interpolated values between each frame. Hence the simulation time step Δ*t* = 2 s is 3 times smaller than the acquisition time, and favors numerical convergence of all time dependent processes. Note that the results were found to be independent of the simulation time step Δ*t* when decreasing it by a factor 2. On the other hand, a segment (or several) that vanishes creates (just before disappearing) an added inflow of 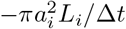, where *a_i_* the segment’s radius just before disappearing. This corresponds to radius retraction as observed in the data.

### 4. Data analysis – Time averages

For all data, we extract short time averages by using a custom-developed MATLAB (The MathWorks) routine. To determine the short time averages of the oscillating shear rate and vein radius, we used a moving average with a window size of *t*_ave_ ≃ 2 – 3T (*T* ≃ 120s). The *i*^th^ element of the smoothed signal is given by 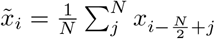, where *N* is the window size. At the boundary where the averaging window and the signal do not overlap completely, a reflected signal was used as compensation. This can be done because the averaging window is relatively small and the average varies slowly in time. The determined trend (for the close-up data sets) was then smoothed with a Gaussian kernel to reduce artefacts of the moving average filter.

In experimental data of the shear rate, we observe that raw shear rate appear to oscillate at rather high frequency (see *e.g*. Fig. 1-iii). Here we briefly rationalize this behavior. First a zoom in time of the data in Fig. 1 A iii, see App. 1 - figure 1, shows that in fact the frequency at which raw shear rate oscillates is double that of the frequency of oscillations of the vein radius. We explain this frequency doubling based on a minimal example. Consider a minimal example with a contraction pattern *a*(*t*) ≃ 〈*a*〉(*t*)(1 + *ϵ*cos(2*πt*/*T*)), where the average radius slowly evolves in time as 〈*a*〉 = *L* cos(2*πt*/*t*_adapt_). The flow in the vein is 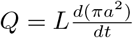 and therefore the shear rate at lowest order in ϵ is

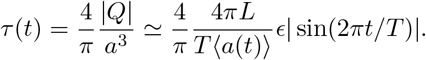

The resulting shear rate contains the absolute value of a periodic quantity of period *T*, hence, is periodic with half the period *T*/2. We plot the minimal example curves in App. 1 - figure 1-B.

We further check whether our algorithm to extract the shear rate trend is correct even on these high frequency raw data. Averaging the raw shear rate obtained in the above minimal model over one contraction period yields

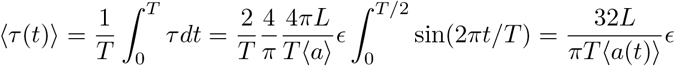

which is exactly the amplitude of the raw *τ* data up to a constant numerical prefactor. Hence, our averaging is well suited to extract reliable trends of the shear rate. In App. 1 - figure 1-B, we present the results from our averaging algorithm (full thick red line) and the theoretically calculated trend (yellow dashed) and obtain excellent agreement. Our time-averaging algorithm is therefore well-suited to the investigation of even these high frequency data.

**Appendix 1 - figure 1.**
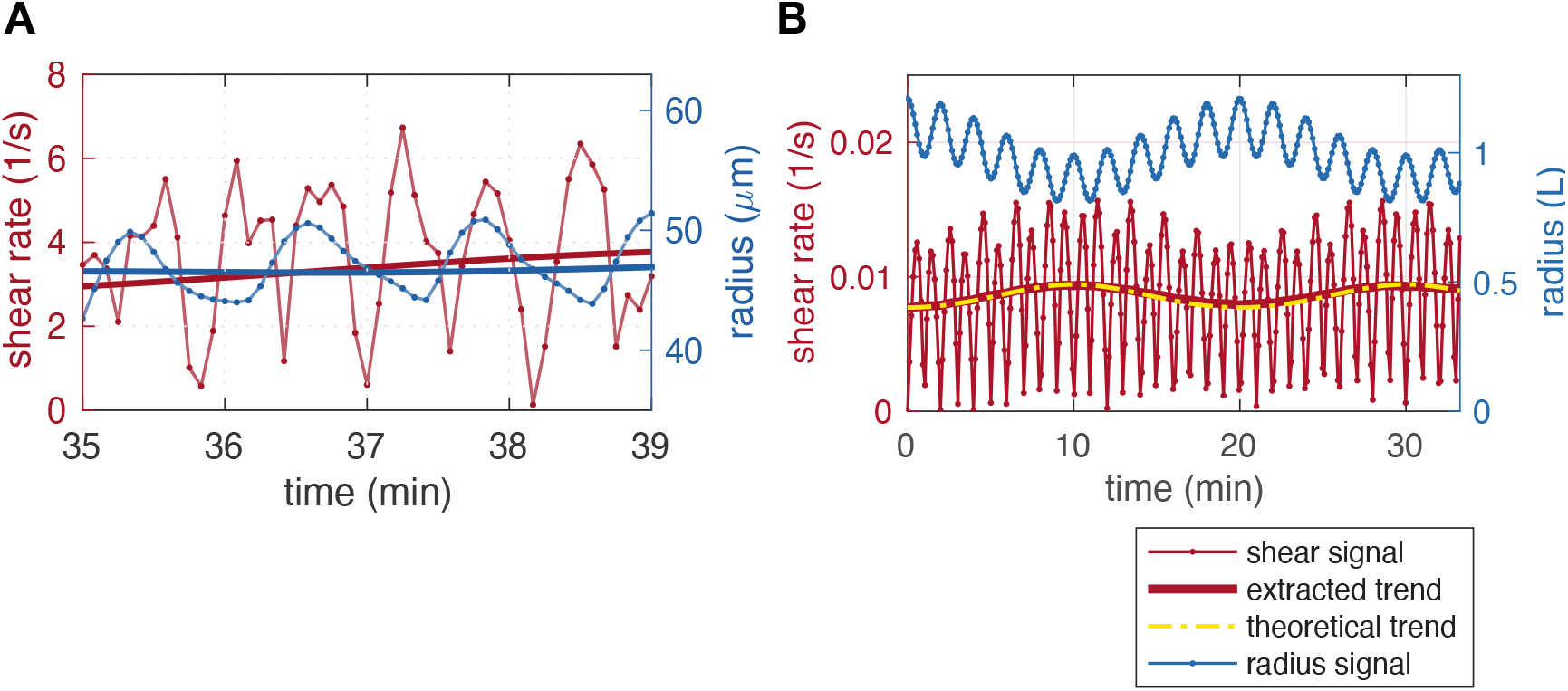
Extracting average shear rates from shear rate data. (A) Time zoom of a close-up data set (that of Fig. 1 A iii, #*K*) showing the doubling of the frequency of the shear rate compared to the radius. (B) minimal model example with short timescale and long timescale radius oscillations, resulting in shear rate with a doubled frequency. Here the contraction period *T* = 2 min and *t*_adapt_ = 20 min.

### 5. Data analysis – Additional shear rate - radius data

To add to the data presented in Fig. 1 iv presenting the time-averaged dynamics of radius adaptation and shear rate, we show in App. 1 - figure 2 (resp. App. 1 - figure 3) additional dynamics for the close-up datasets (respectively the full network #1 of Fig. 1-B).

**Appendix 1 - figure 2.**
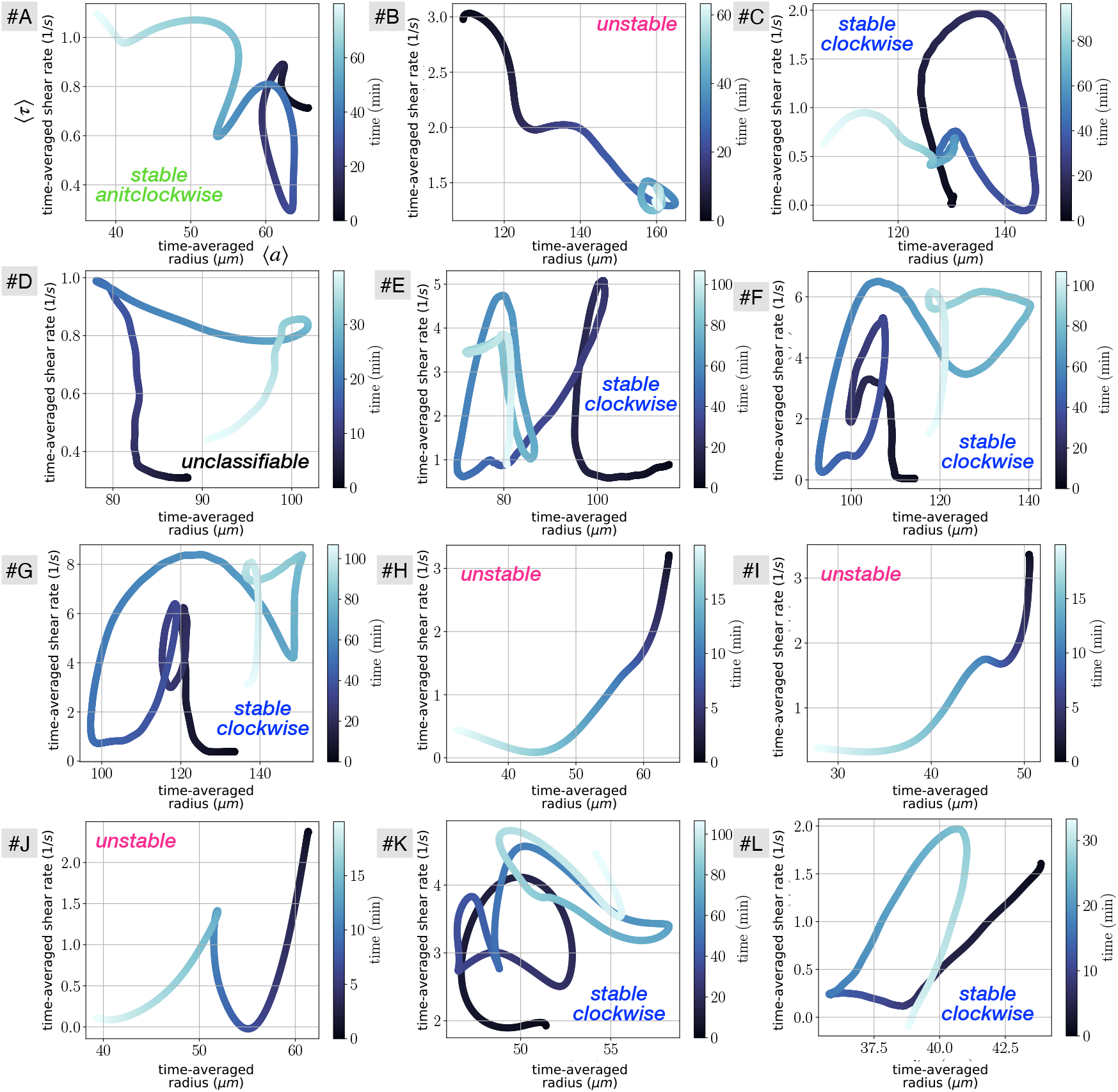
Vascular adaptation dynamics for all close up experiments, using time-averaged shear rates 〈*r*〉 and time-averaged radius 〈*a*〉. The letters indicate the data set names, and are used consistently throughout the manuscript. Typical classification of vein dynamics is indicated for each plot.

**Appendix 1 - figure 3.**
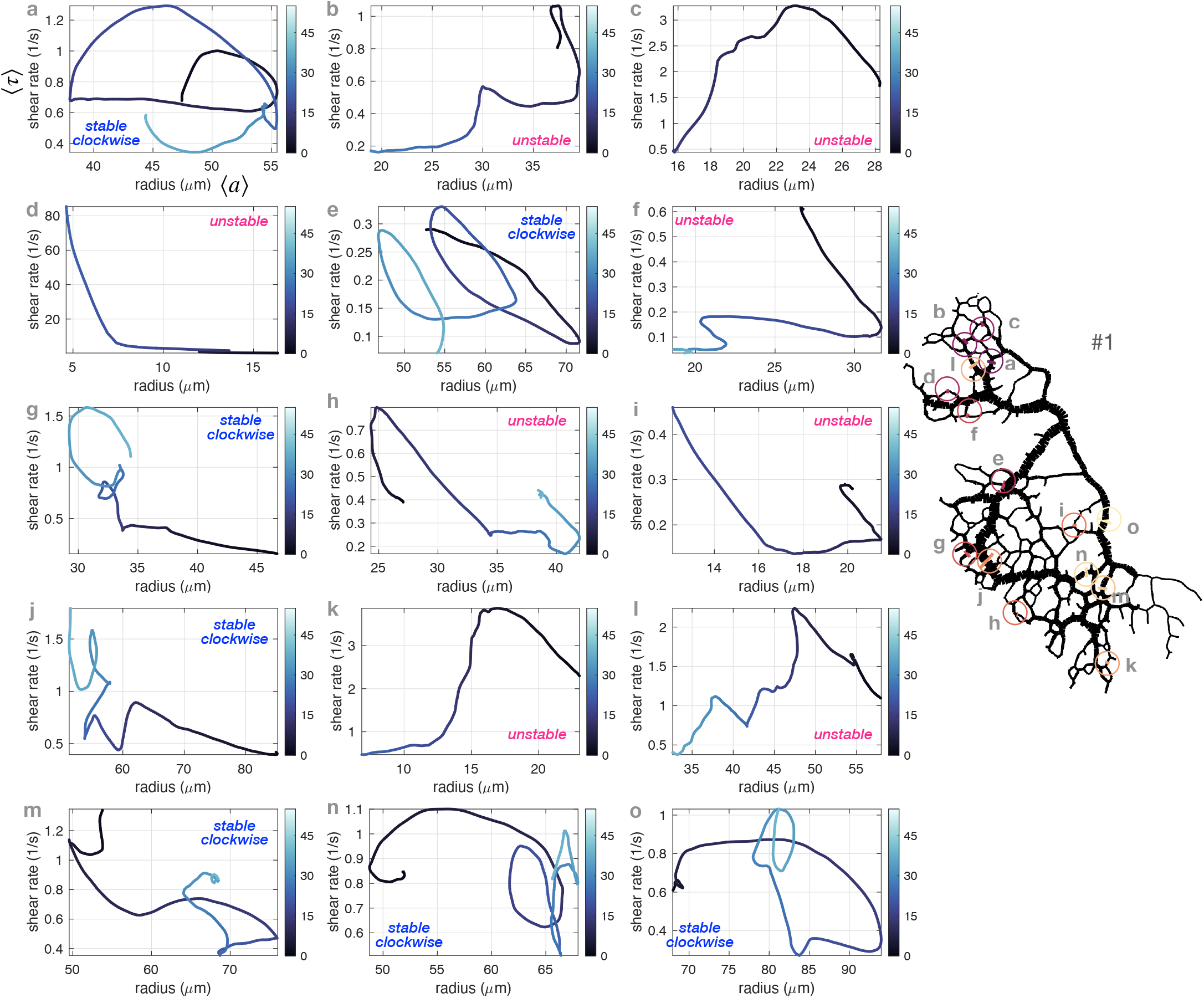
Vascular adaptation dynamics for a few veins in the full network #1 using time-averaged shear rates 〈*τ*〉 and time-averaged radius 〈*a*〉. The veins shown are chosen randomly but distributed throughout the network. The network sketch on the right hand side shows circles indicating at their center the location of each vein, with consistent labels. Typical classification of vein dynamics is indicated for each plot.

### 6. Data analysis – Additional data on full networks

#### a. Additional data on different full network specimen

In what follows we present additional data on full networks. In particular, we investigate two other full networks besides specimen #1 (of Fig. 1-B), which we call #2 and #3. These two additional networks show significant spontaneous reorganization over time and we show snapshots of their initial and final networks in App. 1 - figure 4-A and B and App. 1 - figure 5-A and B.

We also present additional data to demonstrate the existence of similar ambiguity in shear rate - radius response in other full networks. We show with yellow arrows additional places where shear rate is initially high yet the vein will disappear in App. 1 - figure 4-C and App. 1 - figure 5-C. Red dots in App. 1 - figure 4-B and App. 1 - figure 5-B also show veins where shear rate is initially low however these veins will grow in time.

We present pressure data in App. 1 - figure 4-D and App. 1 - figure 5-D. We find that pressure doesn’t vary much throughout the network. A global pressure wave is observed corresponding to a stable direction of the peristaltic contractions. We identify in these maps nearby veins and find that the ones with larger pressure remain (blue stable) while those with lower pressure vanish (pink unstable).

We present relative resistance data *R/R*_net_ in App. 1 - figure 4-E and App. 1 - figure 5-E. We find a number of veins with *R/R*_net_ > 1, indicated by pink arrows, that indeed vanish in time.

To finish with the analysis of additional networks, we present a map of the time of disappearance of veins in the full specimen in App. 1 - figure 4-F and App. 1 - figure 5-F. We find that loops consistently vanish by their center, as highlighted via black arrows.

#### b. Additional data on full network specimen #1

In App. 1 - figure 6 we present additional data on Specimen # 1 that is the main example under scrutiny in the main text. We provide in particular maps of quantities that are not shown in the main text, such as the connected resistance *R*_net_ (C) and 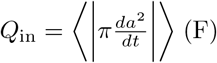. We find that *Q*_in_ typically evolves like the vein radius: showing larger values (light blue) for larger veins and reciprocally smaller values (dark blue) for smaller veins. *R*_net_ in contrast evolves quite dramatically from vein to vein, according to how the vein is close or not to major highways.

We also provide cross-correlation data between specific quantities and initial vein radius 〈*a*〉 at the beginning of the experiment, *R*_net_, *R/R*_net_, *Q*_in_ and 〈*P*〉 (A,B,D,E). We find that the only quantity that is significantly correlated with 〈*a*〉 is *Q*_in_, coherently since we expect *Q*_in_ ∝ 〈*a*^2^〉.

**Appendix 1 - figure 4.**
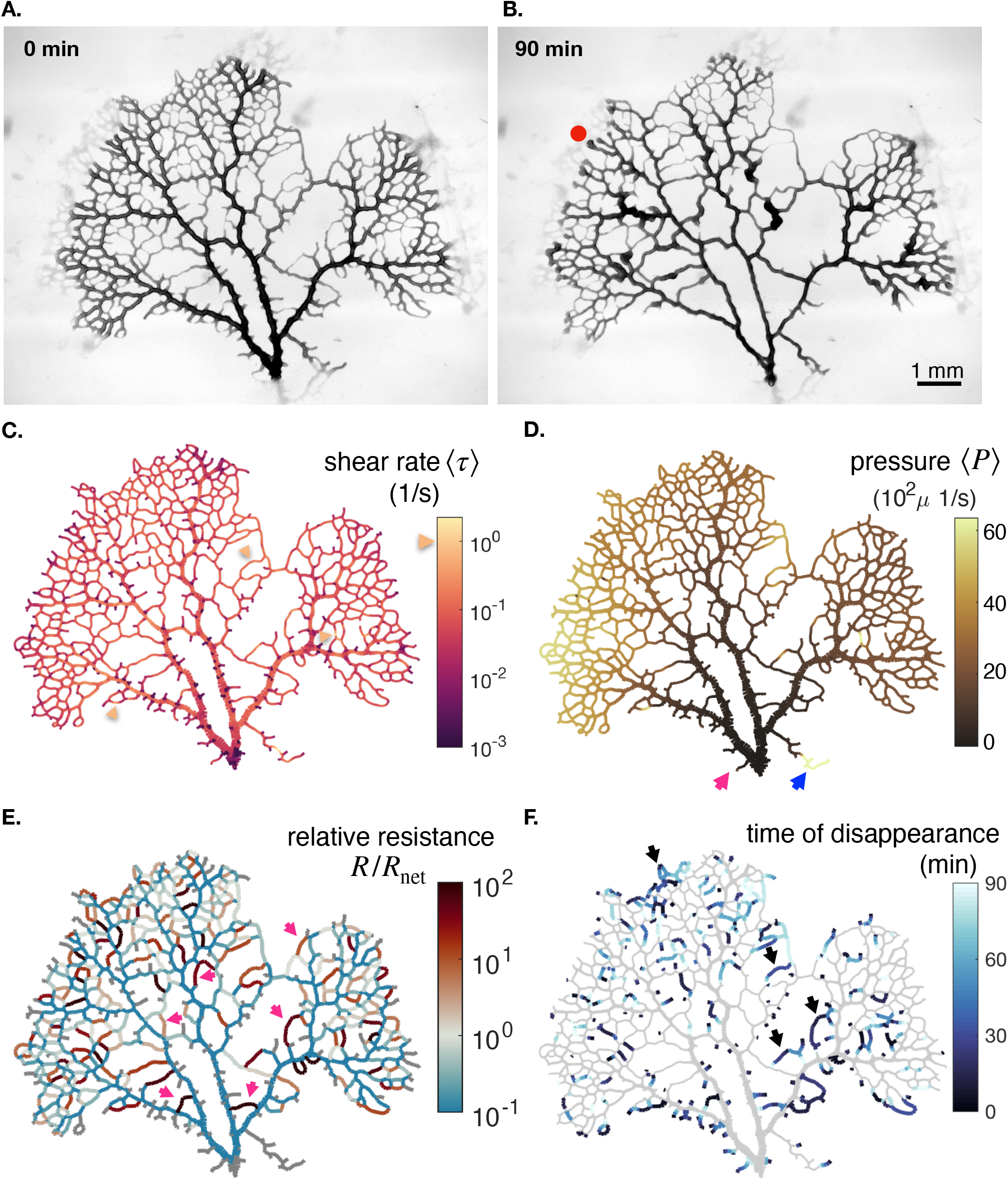
Additional data on the full network specimen #2. (A - B) Bright field images of a specimen with long time dynamics of vanishing veins, specimen 2. (c) Mean shear rate 〈*τ*〉, (D) pressure 〈*P*〉 and (E) relative resistance *R/R*_net_ at the initial stage. (F) Time of vein disappearance for the entire experiment. See text for more information on the arrows and what to take out from the color maps.

**Appendix 1 - figure 5.**
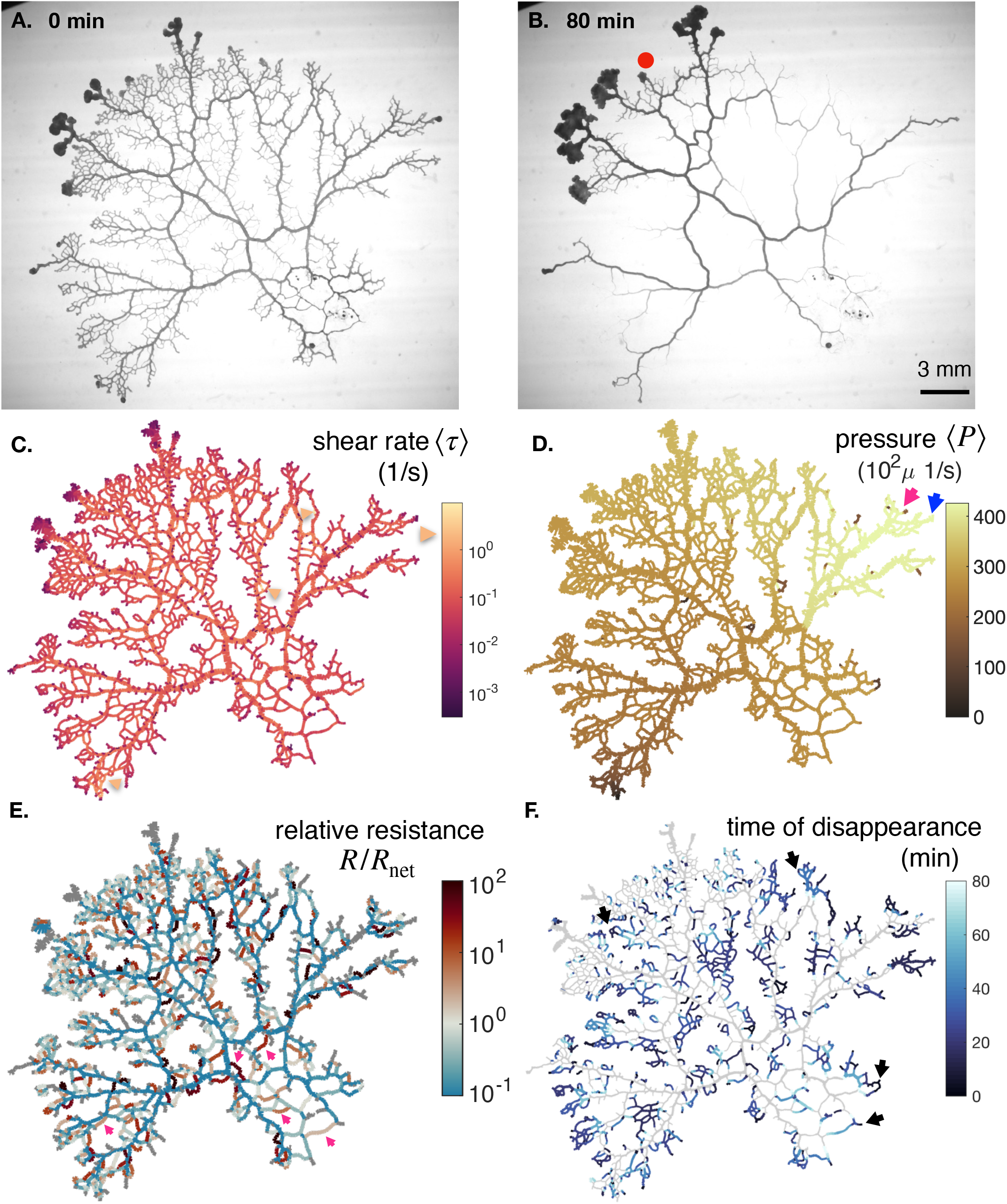
Additional data on the full network specimen #3. (A - B) Bright field images of a specimen with long time dynamics of vanishing veins, specimen 3. (c) Mean shear rate 〈*τ*〉, (D) pressure 〈*P*〉 and (E) relative resistance *R/R*_net_ at the initial stage. (F) Time of vein disappearance for the entire experiment. See text for more information on the arrows and what to take out from the color maps.

**Appendix 1 - figure 6.**
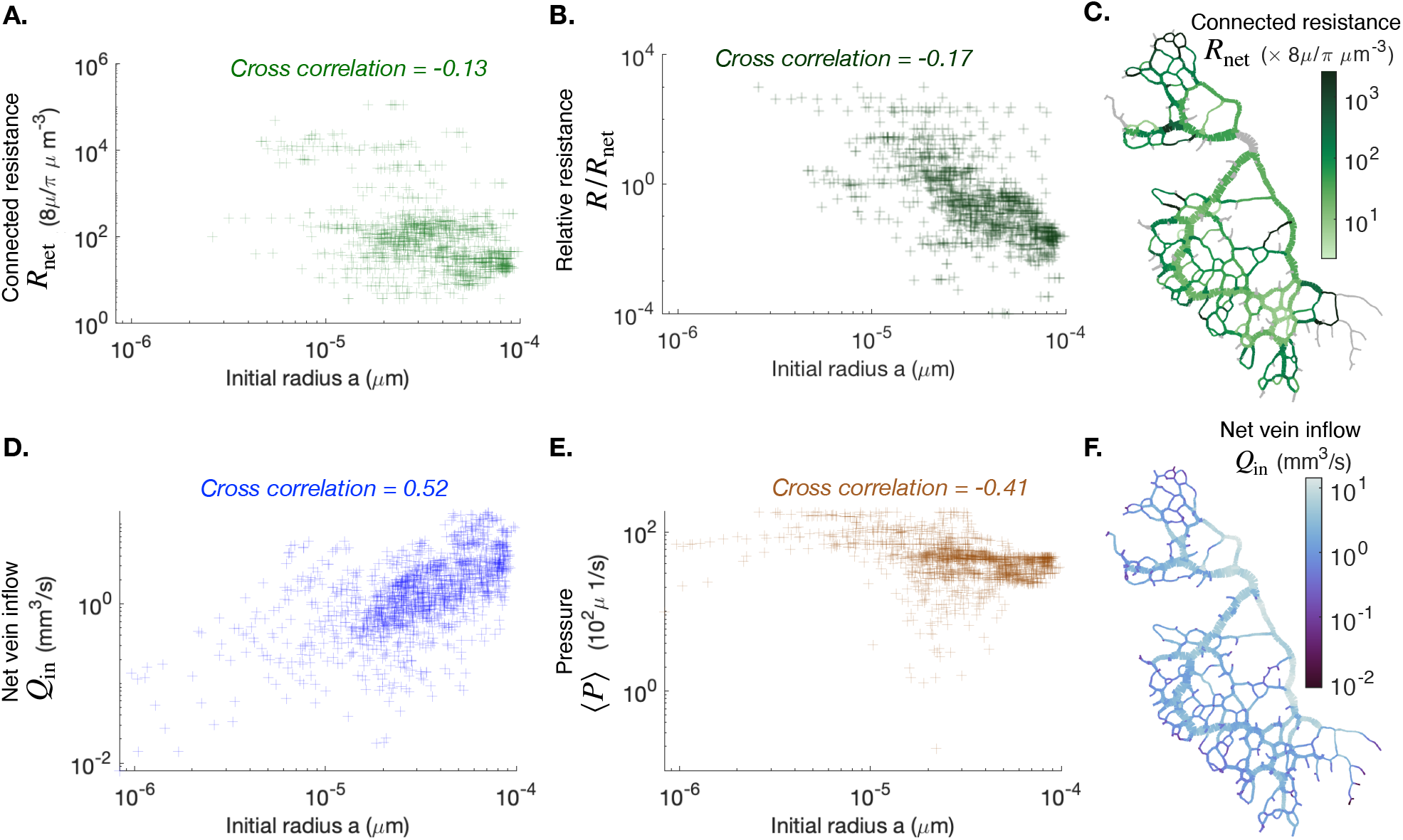
Cross correlation between average vein radius and different flow-based parameters (A) the connected resistance Rnet, (B) the relative resistance *R/R*_net_, (C) the vein outflow *Q* and (E) the local pressure 〈*P*〉. We also present maps of the connected resistance Rnet in (C) and of the vein outflow *Q* in (F). The color scales indicate the magnitude of each variable in each colored vein. All cross-correlations and maps are done at the initial observation time for the full network specimen #1.

## Appendix 2: Adaptation model, time delay and fitting procedure

### 1. Vascular adaptation from force balance

We briefly summarize here the derivation of our vascular adaptation model from force balance and provide more details in our accompanying publication [43]. We consider the force balance equation on a small vein wall segment of radius a, length L, thickness e. As the wall motion is typically slow and occurring over microscopic scales we neglect inertial contributions and write

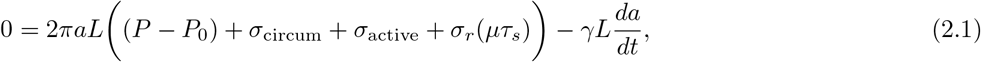

where *P* – *P*_0_ is the hydrodynamic pressure difference between interior and exterior, *σ*_circum_ is the circumferential stress (or elastic tension), *σ*_active_ corresponds to active stresses from the actomyosin cortex [59, 60], and 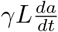 is the friction force reflecting the long timescale for fiber rearrangement [36, 37]. Note that since the shear rate *τ* acts longitudinally on the walls, it does not contribute to the force balance on the radial direction. Yet, the vein walls consist of a material with an anisotropic response to shear, namely cross-linked fibers (the actomyosin gel). Hence, when sheared, a radial stress *σ_r_* (*μτ_s_*) builds up as a result of longitudinal shear rate sensing (with a time delay) [44–46, 61–63].

The general force balance (2.1) significantly simplifies when we average over the short timescales of vein contractions (1 – 2 min) [35], typically corresponding to elastic deformations, to focus on the longer timescales of 10 — 60 min corresponding to vein wall assembly or disassembly inherited from *e.g*. actin fiber rearrangements [36, 37].

On these longer timescales, significant morphological vein adaptation of 〈*a*〉 occurs. 〈*σ*_active_〉 is a constant as it is expected to vary only on short timescales in line with the periodic contractions. Note also that it is a negative stress, that tends to shrink a vein – this reflects the impact of metabolic cost, here induced by vein wall activity. 〈*σ*_circum_〉 ≃ 0 over short timescales, as such forces are intrinsically elastic forces and hence do not pertain long time features. Finally, our numerical calculations of pressures within observed networks show that 〈*P* – *P*_0_〉 depends smoothly on the location within the network, but barely varies in time [8] (Fig. 4-A). We obtain a time-independent, yet position-specific constant 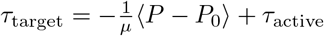, where we wrote *τ*_active_ (*σ*_active_)/*μ*.

Furthermore, we assume a phenomenological functional form for the radial stresses, as 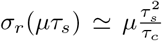, in line with observations of sheared cross-linked actin fibers [44, 45] where *τ_c_* is a positive constant. Importantly, this radial stress, acts in the positive direction, *i.e*. dilates vessels. This functional form is also consistent with measured data on fibrin gels [46, 47] and models of anisotropic response based on nonlinear elastic theory [46].

Finally, to simplify the expressions we now introduce 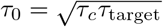 and

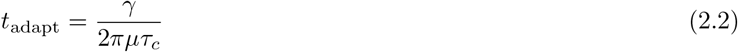

a characteristic adaptation timescale for vascular rearrangement. This allows us to recover the vascular adaptation rule Eq. (1). While the two parameters *τ*_0_ and tadapt may appear to be coupled at the scale of the network, there is actually no reason for *τ_c_* or for *γ* to be constant throughout the network. In fact they may very well depend on the age of the vein, the absolute thickness of the actomyosin gel, *etc*. Again, we refer the reader to more details on the derivation in our accompanying manuscript Ref. [43].

#### Agreement with Murray’s law

Our model is consistent with Murray’s steady state assumption. In fact, the (non-trivial) steady state of our model Eqs. (1–2) corresponds to a constant average shear in the vein 〈*τ*〉 = *τ*_0_. This corresponds exactly to Murray’s result of minimum work.

In fact, Murray stipulates that the energy dissipation 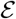 of a single vein (of radius *a* and length *L*) is given by flow dissipation associated with the vein’s resistance and energy expense to sustain the vein

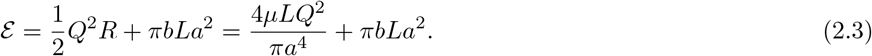

where *R* = 8*μL/πa*^4^ is the vein resistance assuming Poiseuille flow in the vein, *b* is a local metabolic constant per unit volume, *Q* the flow rate and *μ* viscosity. The principle of minimum energy expense suggests to search for the minimum of 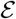 with respect to the vein radius *a* which gives the relation 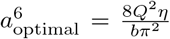. The shear rate *τ* can be expressed as 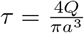 and hence the optimal (or steady state) shear rate is independent of radius and flow rate 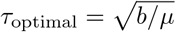. This is consistent with our steady state where shear rate is constant 〈*τ*〉 = *τ*_0_. The constant *τ*_0_ can thus also be interpreted as being related to the typical local energy expense to sustain the vein 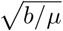 (which corresponds very closely to our *τ*_active_ characterizing metabolic expense to sustain the contractile activity). Note that we bring further insight compared with Murray’s derivation, as our adaptation dynamics (2.1) originates from force balance on the vein wall, and hints that *τ*_0_ (or the metabolic cost) also depends on local pressure 〈*P*〉.

### 2. Extracting the time delay from data analysis

In this section we discuss our procedure to extract the time delay from data.

First, we verify that the time delay we extract is independent of the averaging technique. To do so we investigate the time delays obtained from the cross-correlation of *da/dt* and *τ* instead of their averaged counterparts *d*〈*a*〉/*dt* and 〈*τ*〉. We obtain a distribution of best time delays, over the nearly 10000 vein segments of the full network, and we retain maxima regardless of the value of the cross-correlation. We present the results in App. 2 - figure 1-A. The average time delay is 52 s, which is comparable in orders of magnitude to the average time delay of 122 s for the same network data but where radii and shear rate trends were extracted (Fig. 2-C). Note, that the correlation however is much less clear without extracting trends and in average the correlation score is 0.25 with only 5% of veins achieving a score > 0.2 compared to 0.66 average score with trends with 15% of veins achieving a score > 0.5. Note that the average correlation is quite low because in general the data are not perfectly periodic and smooth. Hence, we decide to keep the analysis on the data trends, that appears to be much more precise.

Second, we check that even if the time delay between adaptation and shear rate is close to the peristaltic contraction frequency (*T* ≃ 1 – 2 min), we are still able to extract it with our method reliably. To do so, we consider model data 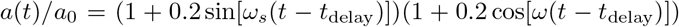 and 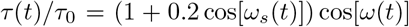. We impose a contraction period *T* = 2*π/ω* = 120 s and the long time adaptation period 2*π/ω_s_* = 20 min, and a delay similar to the beating period *t*_delay_ = *T* = 120 s. Using our methodology to extract the time delay, we find *t*_delay_ = 114 s, which is equal to the set time delay of 120 s within the error bar of 6 s corresponding to the time step where data was sampled. We conclude that the time delay we obtain is independent of the value of the contraction frequency.

Finally, some trajectories appear to oscillate on long timescales say with period *T*_osc_. Hence, it may not be obvious by cross-correlation for these specific trajectories to determine whether the delay is *t*_delay_ or – *T*_osc_ + *t*_delay_, or another combination. *T*_osc_ characterizes rarely observed long cycles in the long time adaptation dynamics, for example see Fig. 2D, and typically *T*_osc_ = 20 min. In contrast, the apparent phase lag between 〈*τ*〉 and 〈*a*〉 is usually of the order of a few minutes in the samples where the delay can be inferred unambiguously (*t*_delay_ ~ 1 – 5min). We may thus expect that the time delay is indeed *t*_delay_ ~ 1 – 5min and not –*T*_osc_ + *t*_delay_ which would be much longer. We impose this condition by adding a cutoff on the time delay at 5 min. Changes to the time delay cutoff, for example setting the cutoff to 10 min, does not affect the results significantly. In fact, strictly oscillatory signals are very rare. For example Fig. 1 B iii clearly shows a lag time (between 7 – 15 min) that allows one to resolve the causality relation unambiguously.

**Appendix 2 - figure 1.**
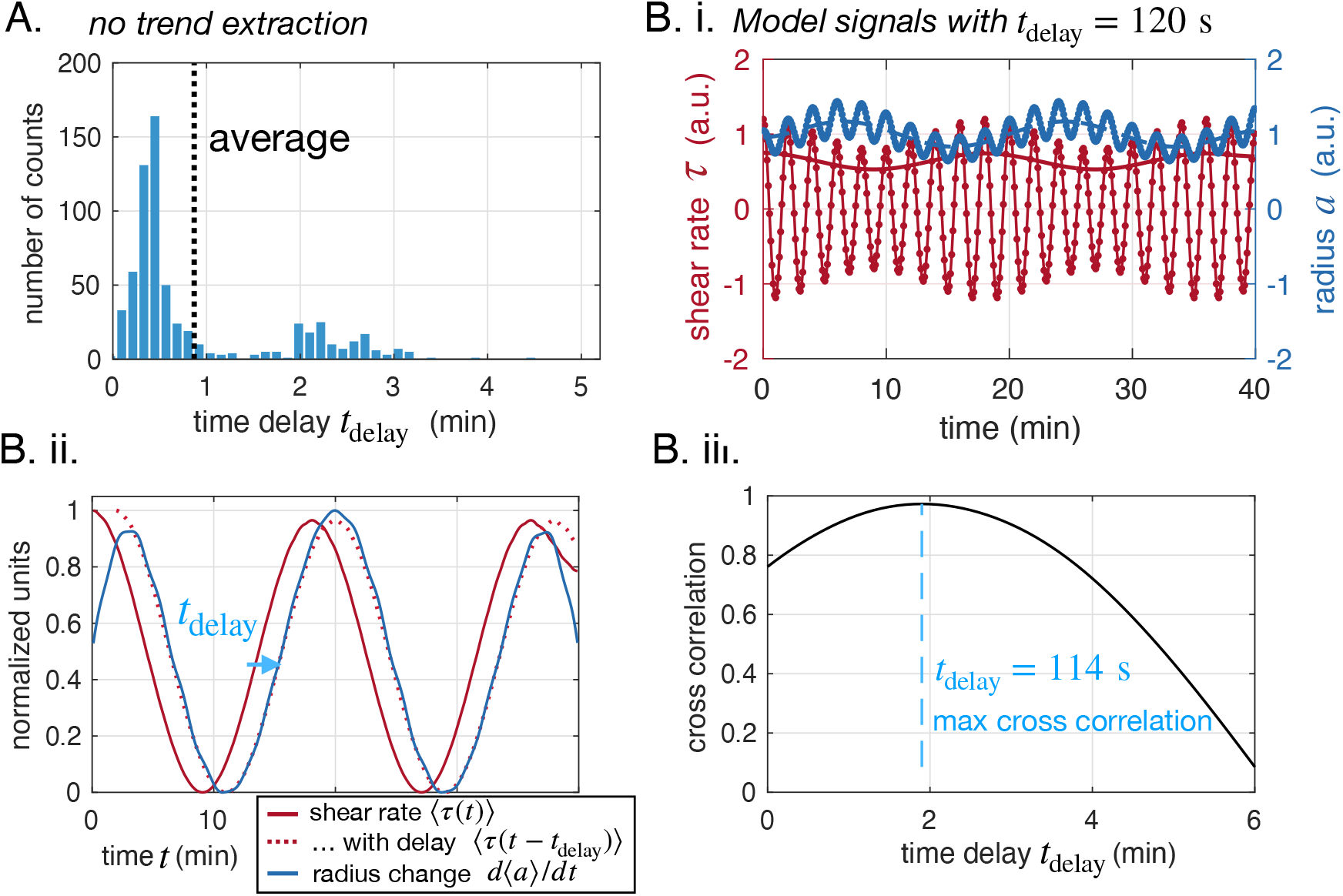
Time delay extracted is independent of averaging technique or oscillation frequency. (A) Time delays measured without extracting trends from data (same plot as Fig. 2-C but without extracting trends). (B) Extracting the time delay for model data with (i) model data and extracted trends, (ii) extracted trends and best time delay obtained shown on trends and (iii) cross correlation between *da/dt*(*t*) and *τ*(*t* – *t*_delay_) with respect to the searched time delay tdelay, and maximum value shown.

### 3. Time delays in close-up and full networks – additional data

We now present time delay analysis in all our specimens.

In App. 2 - figure 2 we present time delay data in full networks. Time delays (both positive and negative) were retained for veins for which the maximum cross-correlation was higher than 0.5. Time delays may only be extracted with sufficient accuracy for stable veins, which do not represent the majority of veins in the network. Hence approximately 15 — 25% of observed veins reach a significant cross-correlation and allow us to record a value of the time delay. To avoid biasing the statistical search with either positive or negative time delays, we allow the algorithm to record both positive and negative time delays for a single vein if these maxima are significant. The phenomenon of negative time delays is quite infrequent. For example in specimen #1, out of the observed veins that yield a time delay, we find 94% with a positive time delay, 4% with a negative time delay, and 2% with both a positive and negative time delay. For specimen #2 we find 96% positive, 3% negative and 1% positive and negative, and for specimen #3 respectively 87%, 12% and 1%. Hence, positive time delays are much more likely. The average time delay is consistently *t*_delay_ ~ 2 min.

**Appendix 2 - figure 2.**
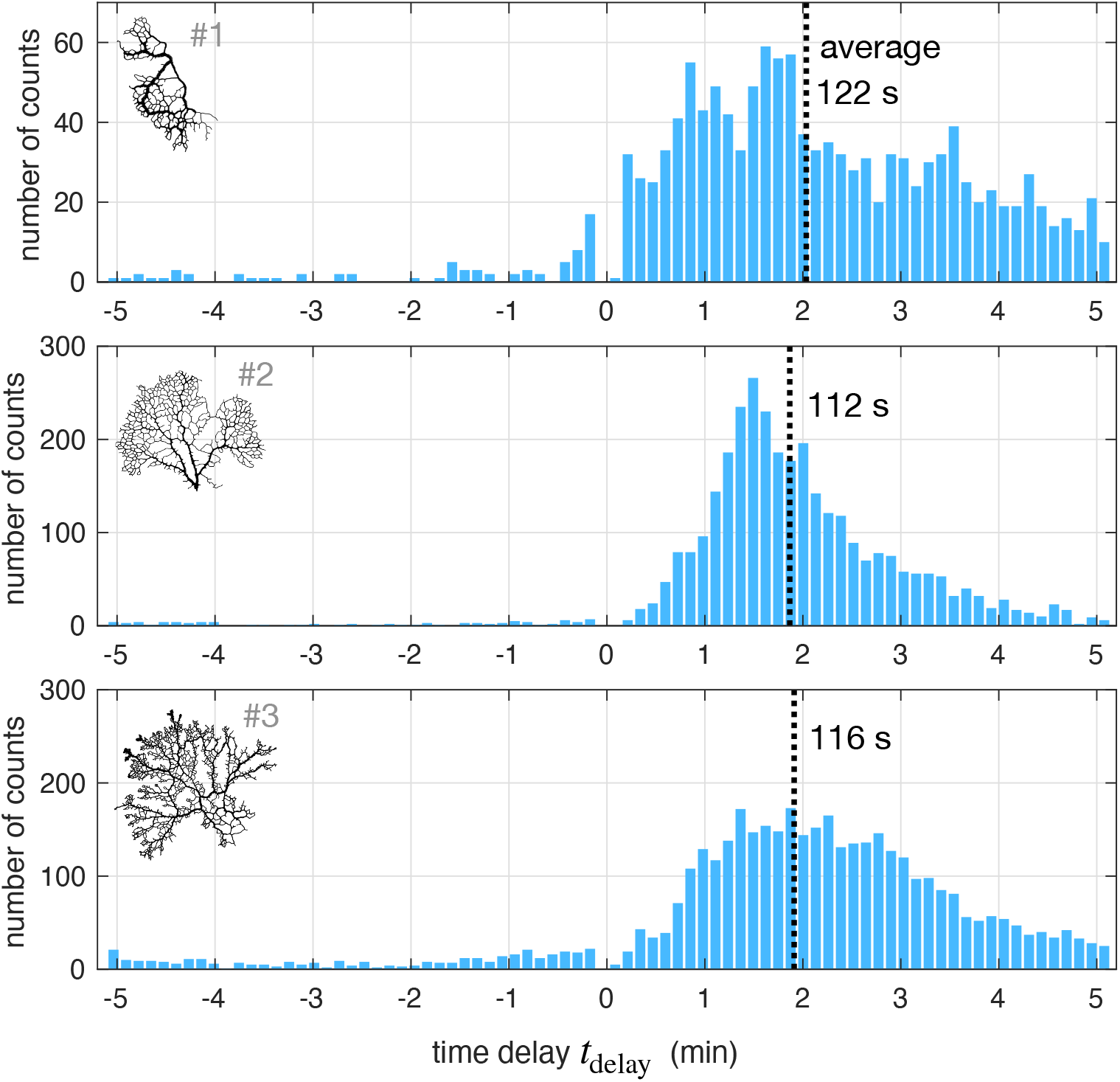
Time delay statistics from full networks. Distribution of best time delays for all veins in the network (#1, with about 10000 vein segments and #2 and #3, both of which have about 30000 vein segments). (insets). Network maps – not to scale.

We also investigate the time delay on close-up data sets (see App. 2 - figure 3), and only on stable close-up data sets as they will allow us to extract the time delay more reliably. Notice that cross correlations are usually quite smooth as the correlation continuously increases until significant shear rate and radius changes are aligned. The correlation maximum corresponds to a strongly correlated configuration (> 0.7). Vein #E finds a best time delay that is quite large (*t*_delay_ ~ 12 min), potentially due to the fact that we are exploring a very long time sequence for this particular vein and that the cross correlation algorithm picks up a large change unrelated to the actual short delay. Notice, however, that a time delay of 2 — 5 min potentially corresponding to the cross-correlation shoulder also seems suited here. The variability in the time delay extracted on close-up data sets show the need for statistical analysis of the time delay, which we perform on full networks.

**Appendix 2 - figure 3.**
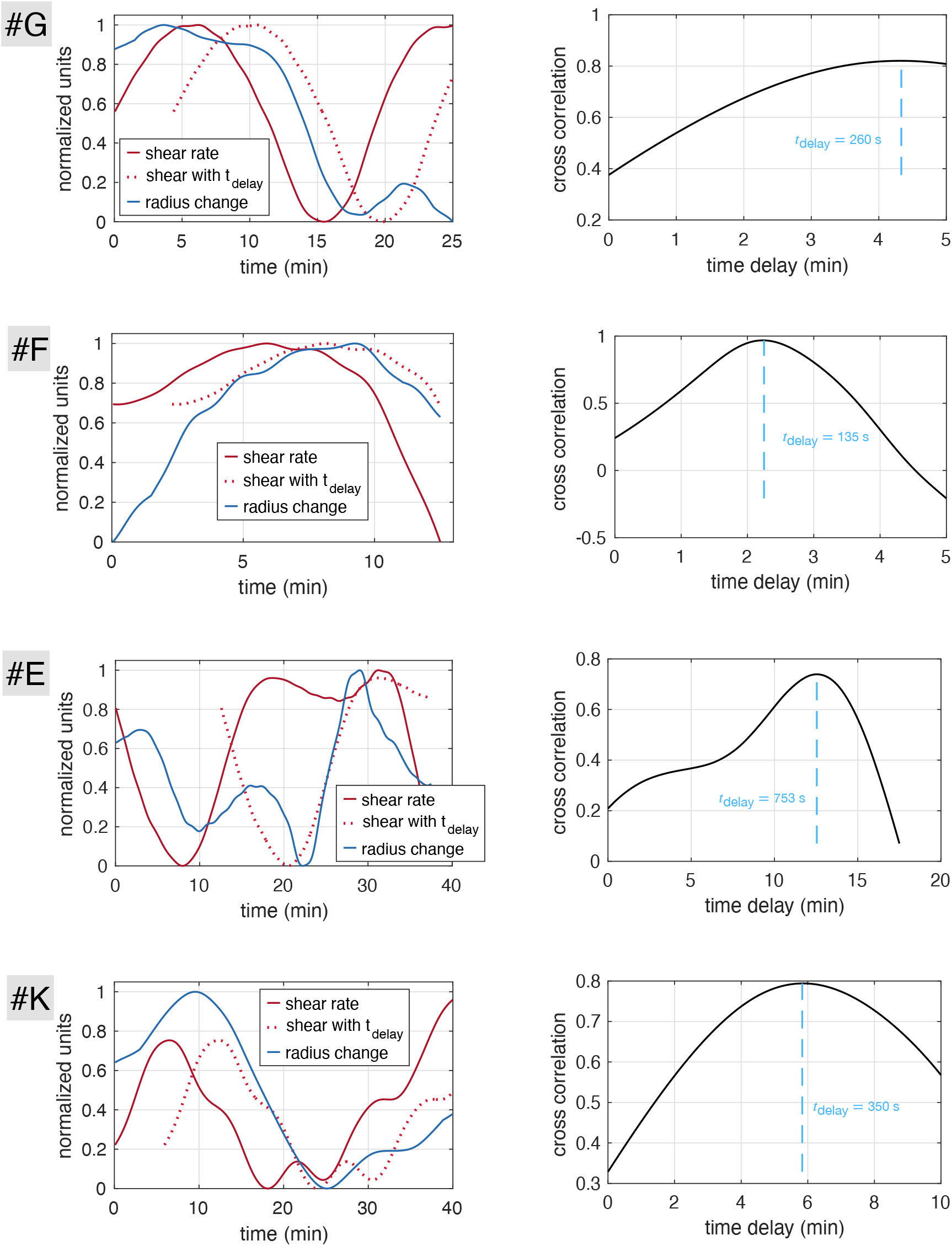
Time delays obtained with cross-correlation method on stable close-up data sets. (Left hand side) Time-averaged shear rate 〈*τ*〉 (red) and radius change (*d*〈*a*〉/*dt*) with time for each vein (#E, #F, #G, #K), as well as time delayed shear right producing the best cross correlation 〈*τ*(*t* – *t*_delay_)〉. (Right hand side) Cross-correlation with varying time delay and optimal time delay obtained at the correlation maximum.

### 4. Fitting of the model to data

Fitting of the model Eqs. (1) and (2) to the data was performed using a non-linear least squares algorithm included in the SciPy optimize package [79], or a linear least squares algorithm, according to whether two or three model parameters had to be fitted. The relative fitting error is defined as

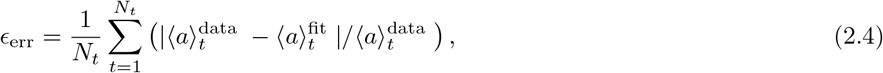

where *N_t_* is the number of data points.

First, we fit close-up data sets, for all three parameters *t*_adapt_, *t*_delay_ and *τ*_0_. As stressed in the main text the model parameters are not expected to be constant over long times (on which loopy trajectories are typically observable). To find suitable time frames where model parameters where approximately constant and loopy trajectories observable we systematically varied the time windows of the data used for fitting. To find the optimal time windows for fitting including fitting the time delay *t*_delay_, we chose close-up data sets forming loopy trajectories (#*G*, #*E*, #*F* and #*K*), as loops are a characteristic feature ensuing from the time delayed dynamics. The distribution of time delays fitted for different time windows was found to range from 1 min to 10 min (see App. 2 - figure 4), which is within the range obtained via the cross-correlation algorithm described above in Appendix 2 3. Fitted trajectories reproduce the main features observed experimentally in detail. The corresponding fitted parameters are reported in Table. 2.1.

Second, we fit all close-up data sets now only including two model parameters, *τ*_0_ and *t*_adapt_. We fixed the time delay to a constant value *t*_delay_ = 120s. We fit different time intervals in the data sets and find very good agreement between data and fits - see App. 2 - figure 5. We report the corresponding fitted parameters in Table. 2.2.

Finally, we fit a random sample of 15 veins from the full network specimen #1. We include two model parameters, *τ*_0_ and *t*_adapt_. We fixed the time delay to the value obtained by cross correlation. We fit only over one rather larger time interval of about 40 min and find reasonable agreement between data and fit (see App. 2 - figure 6). The corresponding fitted parameters are reported in Table. 2.3. In addition for the 15 veins from the full network we also fit the model with no time delay *t*_delay_ = 0. For these fits, we set *τ_s_* = 〈*τ*〉 instead of Eq. (2). We show the fitted results in black dotted lines in App. 2 - figure 6 and report here only the corresponding fitting error *ϵ*_err_ in Table. 2.3. We find a systematic higher fitting error for fits without time delay over those with time delay.

**Table 2.1.** Summary of all fitting parameters of sample trajectories depicted in App. 2 - figure 4, right-hand side. These fits include fitting of *t*_delay_). The fits are done over a range in time *t* ∈ [*t*_min_,*t*_max_].

| Experiment | $t_{\text{delay}}$ in s | $t_{\text{adapt}}$ in s | $\tau_0$ in $\text{s}^{-1}$ | $t_{\text{min}} - t_{\text{max}}$ in min | error $\epsilon_{\text{err}}$ |
| --- | --- | --- | --- | --- | --- |
| data set G* | $69 \pm 3$ | $903 \pm 190$ | $5.4 \pm 0.04$ | 25 – 46 | 1.6 % |
| data set F* | $82 \pm 3$ | $335 \pm 8$ | $5.7 \pm 0.03$ | 90 – 102 | 0.17 % |
| data set E* | $352 \pm 7$ | $798 \pm 3$ | $3.07 \pm 0.02$ | 67 – 104 | 0.6 % |
| data set K* | $76 \pm 4$ | $85 \pm 4$ | $3.63 \pm 0.01$ | 50 – 90 | 2.7 % |

**Appendix 2 - figure 4.**
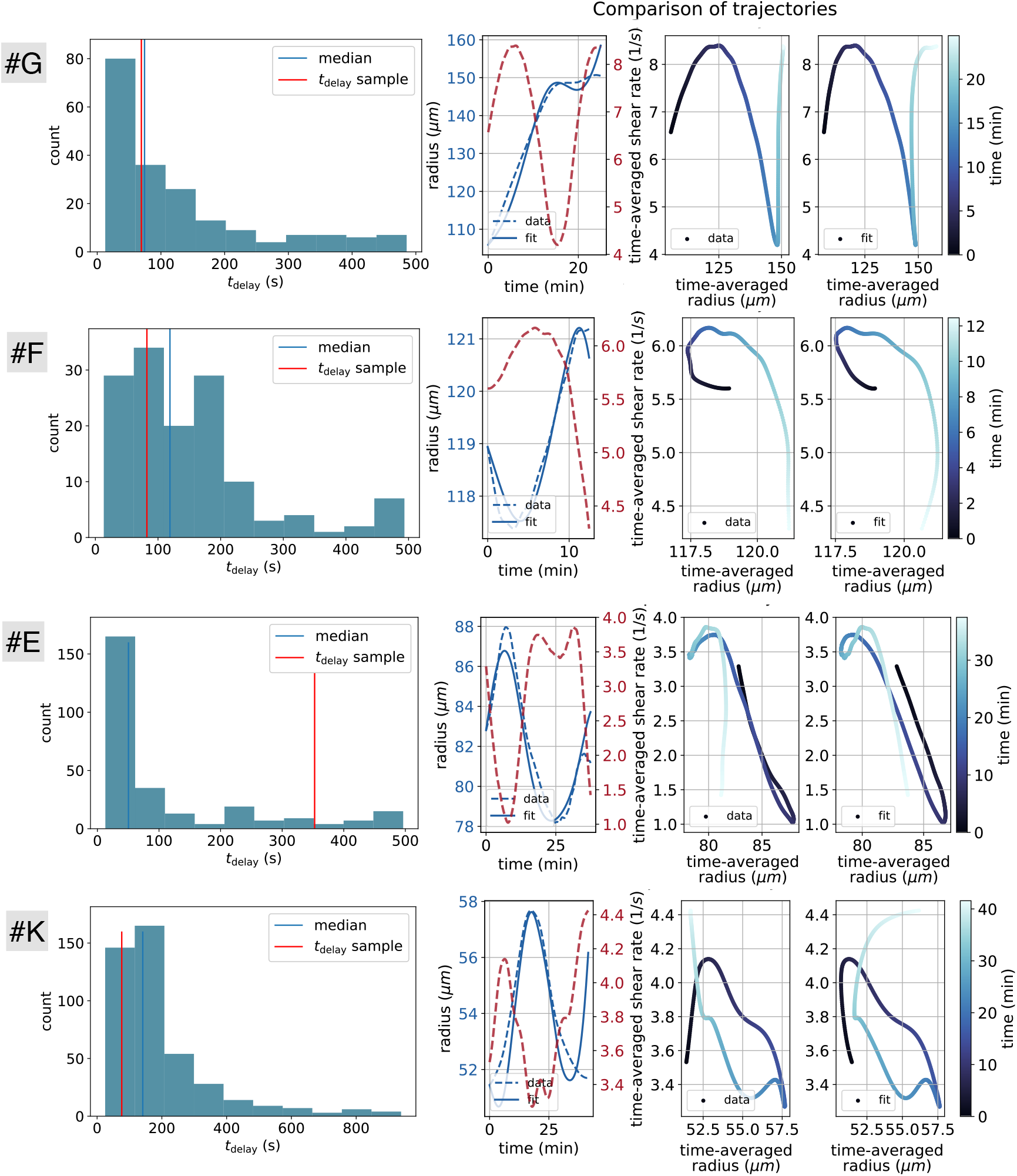
Evaluation of all three model parameters *t*_adapt_, *t*_delay_ and *τ*_0_ from suitable data sets (#E, #F, #G, #K). In the left column the distribution of the obtained time delays using Eq. (1) and Eq. (2) over a distribution of time windows is depicted. To obtain time windows of approximately constant model parameters we performed a fit for every possible time window with the constraints of a reasonable range of fitting parameters and time windows greater than 10 min. The right columns (three graphs) depict a sample of the results of a fitted trajectory, with a given time delay tdelay highlighted on the left hand side graphs as “*t*_delay_ sample”. Among the three graphs on the right, the first, shows the fitted radius data as a function of time and the two next, show the data and the fitted result trajectories in the phase space with shear rate and radius.

**Appendix 2 - figure 5.**
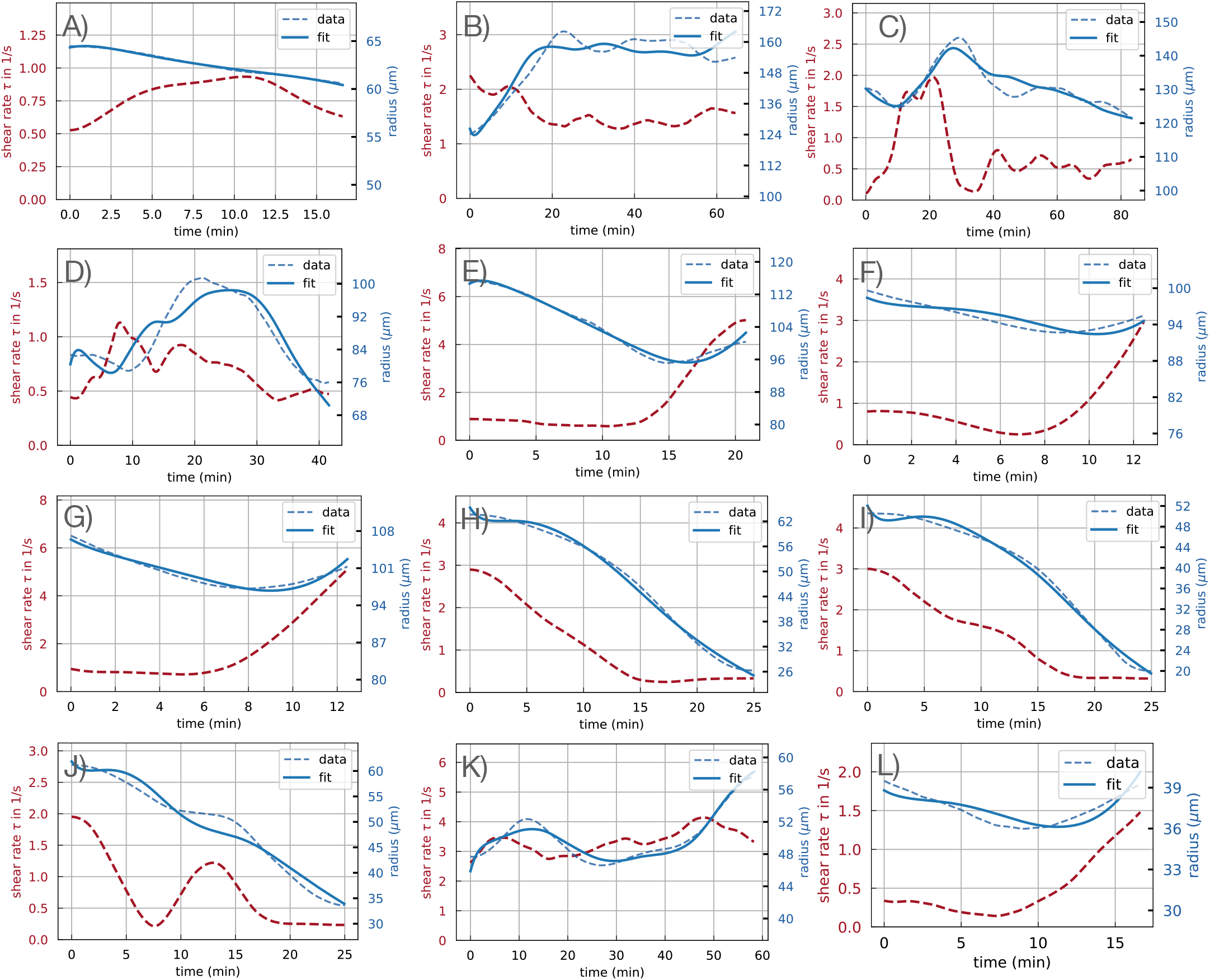
Fit results graphical representation of model Eq. (1) and Eq. (2) using a fixed time delay of *t*_delay_ =120 s for all 12 close-up data sets on a given time window for each data set. The obtained fit parameters are reported in Table. 2.2. All shear rate and radius data presented is time-averaged.

**Table 2.2.** Summary of all fitting parameters with a fixed time delay of *t*_delay_ = 120 s for the 12 close-up data sets. Note that when a data set name is repeated, it corresponds to fitting results over different time ranges *t* ∈ [*t*_min_,*t*_max_] for the same data set.

| Experiment | $t_{\text{adapt}}$ in s | $\tau_{\text{loc}}$ in $\text{s}^{-1}$ | $t_{\text{min}} - t_{\text{max}}$ in min | error $\epsilon_{\text{err}}$ |
| --- | --- | --- | --- | --- |
| data set A | $2682 \pm 831$ | $1.48 \pm 0.17$ | 0 – 17 | 0.13 % |
| data set A | $220 \pm 10$ | $0.44 \pm 0.00$ | 17 – 33 | 0.22 % |
| data set A | $246 \pm 960$ | $1.21 \pm 0.59$ | 67 – 83 | 0.68 % |
| data set B | $1195 \pm 20$ | $1.43 \pm 0.00$ | 10 – 75 | 1.4 % |
| data set C | $5948 \pm 122$ | $0.97 \pm 0.00$ | 0 – 83 | 1.9 % |
| data set D | $1531 \pm 69$ | $0.58 \pm 0.01$ | 0 – 33 | 2.0 % |
| data set D | $377 \pm 6$ | $0.79 \pm 0.00$ | 8 – 50 | 3.2 % |
| data set E | $1312 \pm 37$ | $1.46 \pm 0.01$ | 0 – 21 | 0.93 % |
| data set E | $1894 \pm 49$ | $3.05 \pm 0.02$ | 58 – 83 | 0.66 % |
| data set F | $4323 \pm 319$ | $3.30 \pm 0.06$ | 0 – 42 | 1.9 % |
| data set F | $4323 \pm 319$ | $3.30 \pm 0.06$ | 0 – 42 | 1.9 % |
| data set F | $910 \pm 24$ | $5.08 \pm 0.03$ | 83 – 100 | 0.27 % |
| data set G | $1153 \pm 51$ | $6.07 \pm 0.04$ | 12 – 29 | 0.38 % |
| data set G | $1808 \pm 322$ | $1.00 \pm 0.09$ | 42 – 54 | 0.61 % |
| data set G | $1233 \pm 54$ | $6.68 \pm 0.07$ | 83 – 100 | 0.31 % |
| data set H | $355 \pm 8$ | $2.35 \pm 0.02$ | 0 – 25 | 1.4 % |
| data set I | $274 \pm 5$ | $2.27 \pm 0.01$ | 0 – 25 | 1.6 % |
| data set J | $601 \pm 132$ | $1.41 \pm 0.09$ | 0 – 25 | 3.1 % |
| data set K | $278 \pm 6$ | $3.16 \pm 0.00$ | 8 – 67 | 2.0 % |
| data set K | $85 \pm 2$ | $3.70 \pm 0.00$ | 45 – 75 | 0.6 % |
| data set K | $862 \pm 119$ | $3.40 \pm 0.13$ | 67 – 108 | 1.5 % |
| data set K | $123 \pm 2$ | $4.18 \pm 0.00$ | 108 – 133 | 0.53 % |
| data set L | $2553 \pm 248$ | $0.27 \pm 0.01$ | 6 – 23 | 1.3 % |

**Table 2.3.** Summary of fitted parameters for a 15 randomly selected veins in the full network App. 2 - figure 6. *t*_delay_ was established with cross-correlation, while *t*_adapt_ and *τ*_0_ were obtained through linear least-squares fitting. Error bars correspond to the 95% confidence interval and N.R. corresponds to non relevant points for which the 95% confidence interval yielded error bars as big as the parameters themselves and were, hence, deemed non-relevant.

| Vein index in full network | $t_{\text{delay}}$ in s | $t_{\text{adapt}}$ in s | $\tau_0$ in $\text{s}^{-1}$ | error $\epsilon_{\text{err}}$ | error $\epsilon_{\text{err}}$ with $t_{\text{delay}} = 0$. |
| --- | --- | --- | --- | --- | --- |
| a | 24 | $1800 \pm 250$ | $0.76 \pm 0.02$ | 8.1% | 7.9% |
| b | 78 | $1520 \pm 40$ | $0.89 \pm 0.02$ | 2.6% | 2.9% |
| c | 360 | $1500 \pm 150$ | N.R. | 1.4% | 1.4% |
| d | 120 | $1800 \pm 100$ | N.R. | 17% | 23% |
| e | 120 | $900 \pm 70$ | $0.2 \pm 0.05$ | 7.4% | 11% |
| f | 186 | $2750 \pm 100$ | $0.38 \pm 0.05$ | 5.2% | 6.3% |
| g | 42 | $2450 \pm 50$ | $0.94 \pm 0.03$ | 5.0% | 5.1% |
| h | 246 | N.R. | $0.26 \pm 0.03$ | 8% | 11% |
| i | 186 | $870 \pm 40$ | $0.26 \pm 0.01$ | 3.3% | 6.9% |
| j | 54 | $2200 \pm 100$ | $1.1 \pm 0.2$ | 5.2% | 5.0% |
| k | 288 | $730 \pm 50$ | $4.0 \pm 1.0$ | 4.7% | 6.7% |
| l | 108 | $4050 \pm 200$ | N.R. | 1.1% | 1.2% |
| m | 360 | $10000 \pm 3000$ | $0.62 \pm 0.1$ | 6.1% | 8.0% |
| n | 18 | $2050 \pm 200$ | $0.73 \pm 0.02$ | 3.7% | 3.8% |
| o | 54 | $7300 \pm 3000$ | $0.48 \pm 0.12$ | 6.5% | 6.8% |

**Appendix 2 - figure 6.**
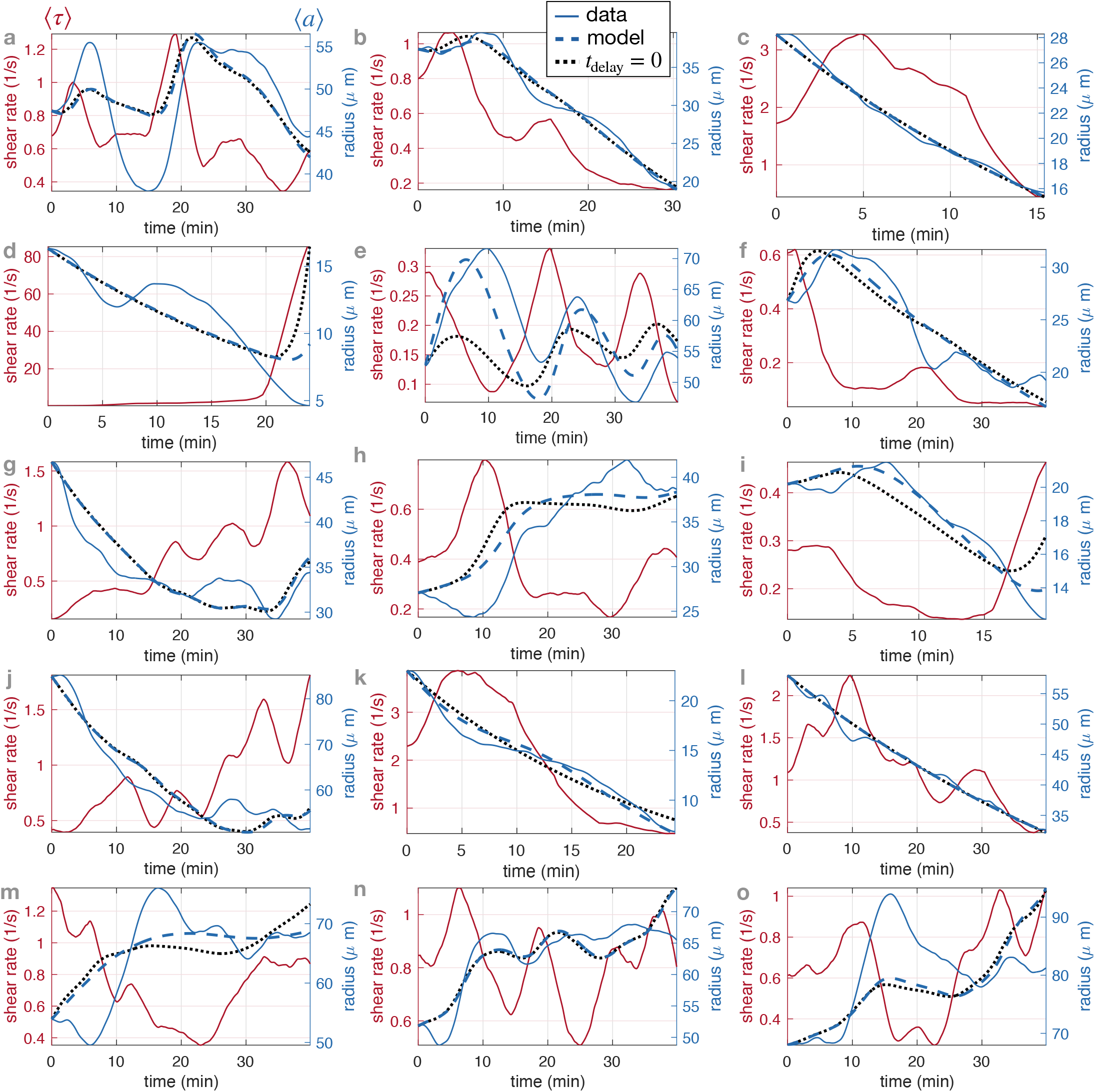
Fit results of model Eq. (1) and Eq. (2) for 15 randomly selected veins on specimen #1. The corresponding fitted parameters are reported in Table. 2.3. The vein positions correspond to those indicated in App. 1 - figure 3.

## Appendix 3: Generic ciruit, stability analysis, and model parameters estimation

### 1. Equivalent resistances

Equivalent resistances (*R*_net_) in our full network structures are calculated using an algorithm based on Kirchhoff’s laws [80], from the values of R for each vein segments directly evaluated from the data-based network architecture. The algorithm was tested to yield correct results on simple geometries where analytic expressions may be found.

We briefly explain the principle of the algorithm and how to interpret the results of *R/R*_net_ on the basis of a few examples in App. 3 - figure 1.

In App. 3 - figure 1(A) a simple network consisting of two veins in series is considered. Considering one of these veins as the vein under scrutiny gives simply that the resistance in the rest of the network is *R*_net_ = *R* since it consists only of one vein. Then *R* = *R*_net_ and the vein is *a priori* stable.

Adding yet another vein in parallel in App. 3 - figure 1(B) modifies the rest of the network. Now it consists in two parallel veins of resistance *R* and hence *R*_net_ = *R*/2 (two resistances in parallel). As a result *R* > *R*_net_ and the vein under scrutiny is *a priori* unstable.

Adding a dangling end to the network App. 3 - figure 1(C) does not modify the resistance of the network attached to the vein. Hence, the vein under scrutiny is still unstable.

**Appendix 3 - figure 1.**
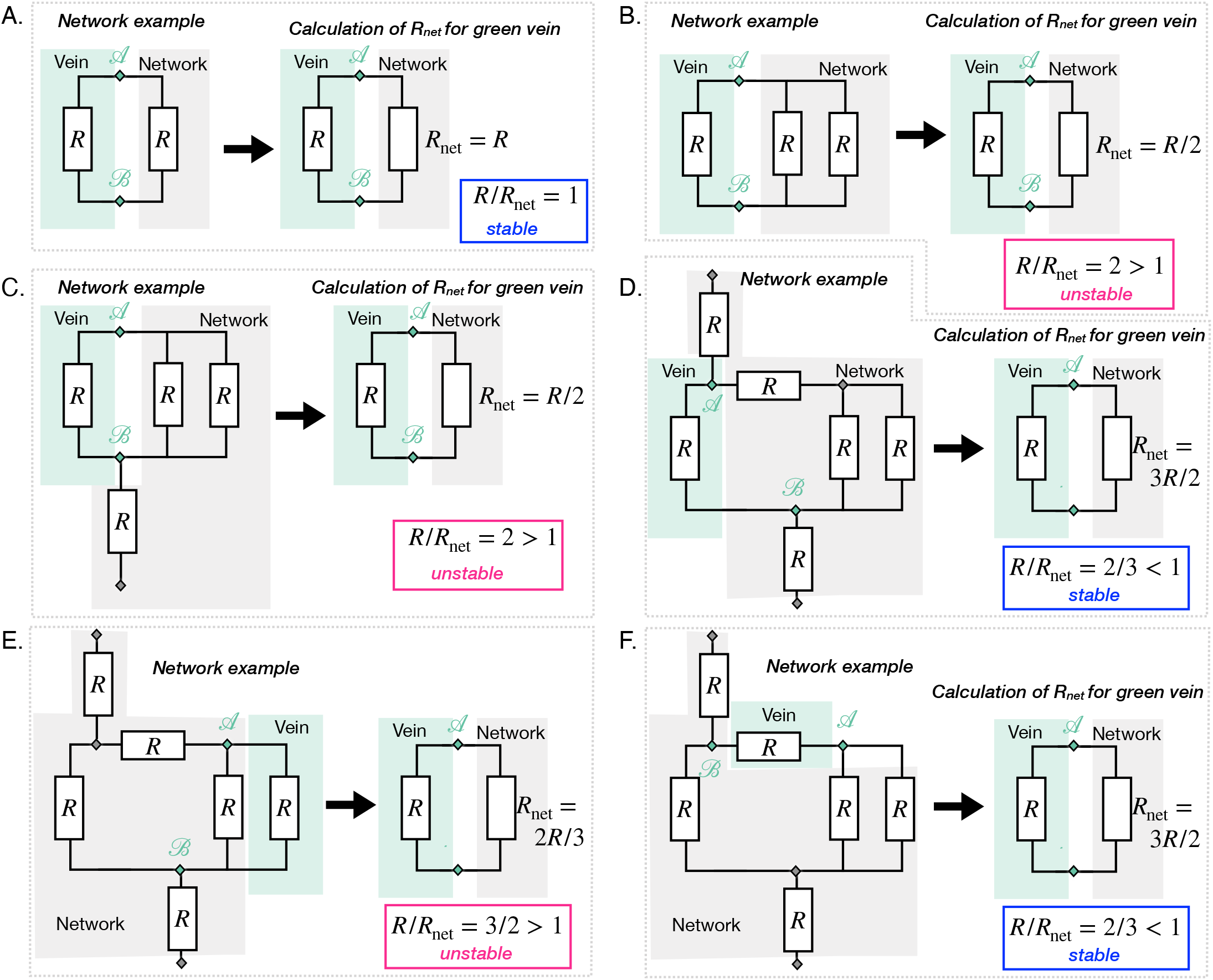
Principle of the calculation of *R*_net_ on the basis of a few examples. The networks are different in all cases except networks D-E-F are the same. D-E-F differ in which vein is under scrutiny. For each case, equivalent resistances *R*_net_ of the rest of the network relative to the vein under scrutiny are calculated via Kirchhoff’s laws. The resulting Rnet is compared to *R*. When *R* > *R*_net_ (respectively *R* < *R*_net_) the vein is unstable, in pink (respectively unstable, in blue).

We make a slightly more complex network in App. 3 - figure 1(D) adding another dangling end and another resistance in series. Again the dangling end does not contribute to the calculation of the equivalent resistance however the vein in series does. We have one vein in series of resistance *R* with two veins in parallel of resistance *R*. The equivalent resistance is *R*_net_ = *R* + *R*/2 = 3*R*/2 > *R* and the vein is *a priori* stable now.

Since this network is slightly more complex we can investigate the fate of other veins in that same network, which we do in App. 3 - figure 1(E-F). We find that these other veins are unstable or stable. This shows that even in a simple network, the relative resistance is a key measure to discriminate between different veins.

### 2. Generic flow network equivalent circuit

We focus on the generic flow network equivalent circuit as given in Fig. 2-A.iii of the main paper and derive the circuit laws as given in the text - see also App. 4 - figure 1-A that recapitulates notations.

Because of Kirchhoff’s laws we easily find that *Q*_in_ = – *Q*_net_. Then we look for the value of the flow rate flowing through the vein of interest *Q*. We see that

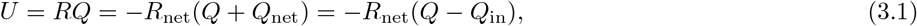

leading to

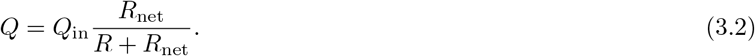

We can then write shear rate in the vein as

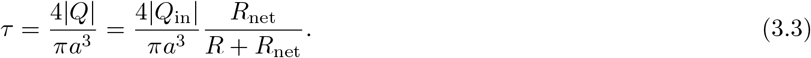

Writing *R* = 8*μL/πa*^4^, *Q* = – *Q*_net_ and averaging over short timescales, we obtain the shear rate at time *t*

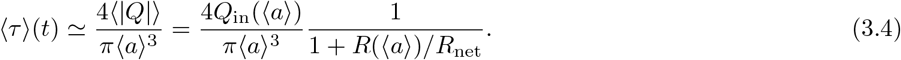

which is exactly Eq. (2) of the main paper.

### 3. Analysis of the feedback system between shear rate and vein radius

In the main text, we have established a set of coupled equations describing the adaptation of veins in a dynamic network. The specific form of these dynamic equations depends on the position of the considered vein within the network. In this section we discuss the stability of a vein fully connected to the network (generic flow network equivalent circuit). Note that other cases (dangling ends, loops, parallel veins) can be easily discussed with similar methodologies.

To simplify the discussion of the fixed points of the dynamical system (〈*τ*〉,*τ_s_*, 〈*a*〉), it is equivalent to study the fixed points of (*τ_s_*, 〈*a*〉), taking into account Eq. (3) in Eq. (2). The dynamic system of equations is then given by:

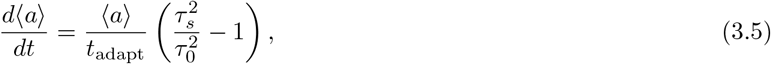

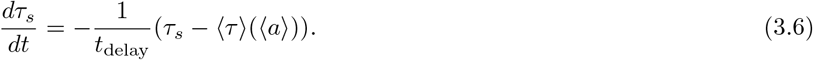

where 〈*τ*〉 is a function of the tube diameter 〈*a*〉:

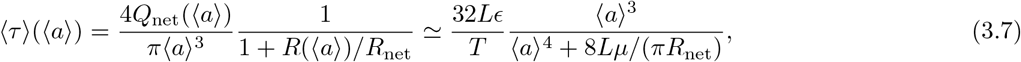

since *Q*_net_ = 8*πLϲ*〈*a*〉^2^/*T* where e is the characteristic contraction percentage of the vein (dimensionless). Plotting the nullclines of Eqs. (3.5) and (3.6) in the (*τ_s_*, 〈*a*〉) space, we observe one, two or no intersections of the nullclines, which correspond to fixed points of the system, depending on the physical parameters. In particular, there is a fixed point corresponding to a vanishing vein in (0, 0). In the following, we will investigate the conditions for the existence and the stability of these fixed points.

#### a. Existence of the fixed points

The dynamical system has more than one fixed point if the nullclines intersect. As depicted in Fig. 2-B, this is the case if max_〈*a*〉_(〈*τ*〉(〈*a*〉)) ≥ *τ*_0_.

The maximum of *τ* (〈*a*〉) is determined by:

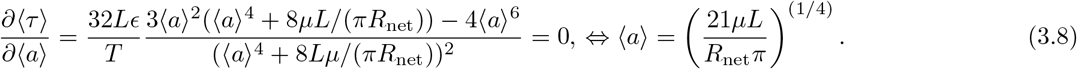

Inserting this in 〈*τ*〉, we get the condition

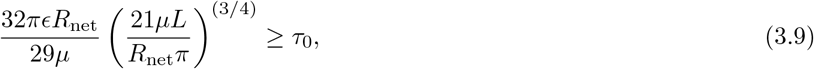

where equality corresponds to one additional fixed point and strict inequality corresponds to two additional fixed points.

#### b. Linear stability of the feedback-system

The dynamical system defined in Eqs. (3.5) and (3.6) has up to three fixed points. To analyze the stability of those fixed points we use linear stability analysis [81, 82]. The first fixed point is at (*τ_s_* = 0, 〈*a*〉 = 0), the other two are defined by (*τ_s_* = *τ*_0_, 〈*a*〉 = *r*_0_, *±*), where *r*_0_, ± are the real positive solutions of the equation *τ*_0_ = 〈*τ*〉(*r*_0_). To analyze the stability of those fixed points, we calculate the Jacobi matrices *J* at each location:

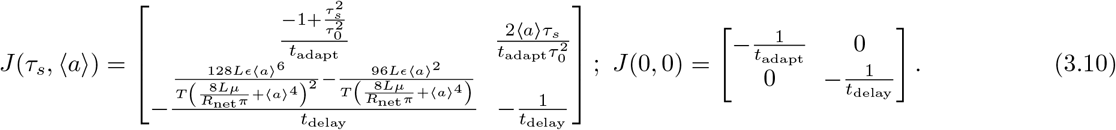

For (0,0) the eigenvalues can be read off from the Jacobi matrix as 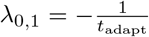 and 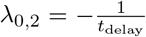. Consequently, the fixed point is stable, as all model parameters are positive. The two other fixed points, as mentioned above depend on the root 〈*a*〉_0_ of *τ*_0_ = 〈*τ*〉(〈*a*〉_0_).

The stability of those fixed points is therefore conditional on the value of these roots. To gain insight on the stability of the fixed points we look at the two extreme cases of either small or large tube radii (as specified below). We will then extend our insight to intermediate tube radii.

- <u>〈*a*〉 → 0</u>: In the case of a small tube radius, we can expand Eq. (3.7) in orders of 〈*a*〉. Expanding up to the first non-trivial order gives:

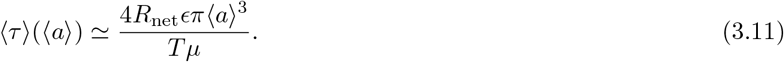 There are thus two fixed points at 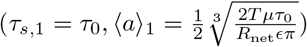 and at (*τ*_*s*,2_ = 0, 〈*a*〉_2_ =0). The resulting Jacobian at (*τ_s_*, 〈*a*〉) is

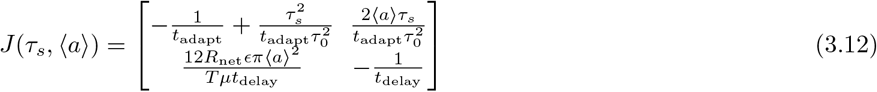 The eigenvalues of *J* at 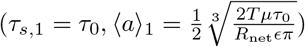 are given by 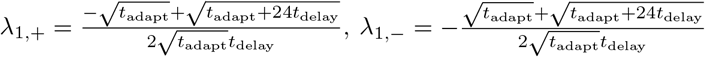. As all model parameters are positive it is easy to see that λ_1,+_ > 0 and λ_1−_ < 0. Consequently, the fixed point is a saddle point. For the second fixed point (0, 0) we recover the same eigenvalues as in the general case, 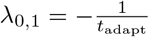 and 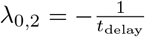, which are both negative and indicate a stable fixed point.
- <u>〈*a*〉 → ∞</u>: In the case of a large tube radius, the shear rate simplifies to 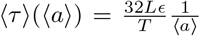 and we find only one fixed point at 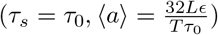. The Jacobian at this fixed point is

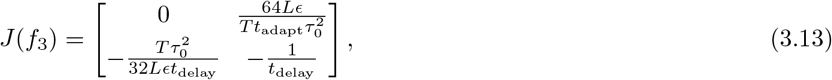

with the eigenvalues

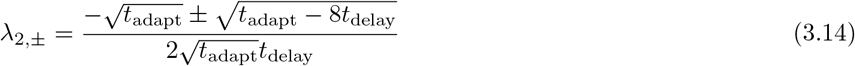 We now have to differentiate two cases. The first one is *t*_adapt_ > 8*t*_delay_. Then *λ*_2,±_ < 0 and the fixed point is stable. For the case *t*_adapt_ < 8*t*_delay_, we have 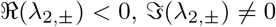 and the fixed point is a stable spiral, which introduces an additional rotation to the system’s trajectories. To investigate the direction of the rotation of this hypothetical spiral, one can look at the sign of Eq. (3.5) for positive displacements *δ* along the shear rate axis. We find that

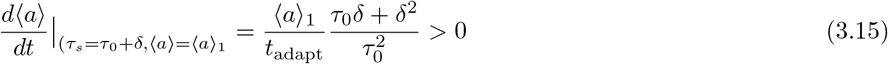

and therefore the spiral rotates in the clockwise direction.

In summary, our stability analysis has shown that the system has up to three fixed points. Two of them are stable and separated by a saddle point. The qualitative stability in the limiting case is also valid for intermediate tube radii, as the stability of a fixed point only changes when two fixed points collide, which is only the case at the bifurcation point (when ∹*a*∪_0_ corresponds to the maximum of *τ*(∩*a*∪)) [81].

## Appendix 4: Equivalent vein flow circuit models for other network topologies of a vein

### 1. Dangling ends

We investigate dangling ends as shown in the network depicted in App. 4 - figure 1-B. Here we consider that a dangling end is connected to the rest of the network at a node where pressure is 〈*P*〉. Since the vein is a dangling end, the only flow flowing through the vein is that generated by peristaltic contractions *Q* = *Q*_in_. Then the shear rate through the dangling vein is simply

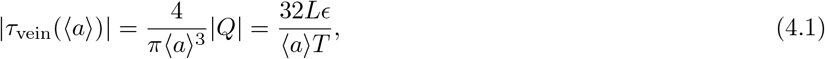

again since *Q* ~ 8*πLe*〈*a*〉 ^2^/*T* where e is the relative contraction amplitude of the vein. We see that this shear rate is the same as in the limiting case 〈*a*〉 → ∞ for the generic circuit. Consequently, this expression does not give rise to any stable non-zero fixed point. Hence, the vein either vanishes or grows indefinitely - see App. 4 - figure 1-B. The crossover between the two regimes occurs for a critical radius

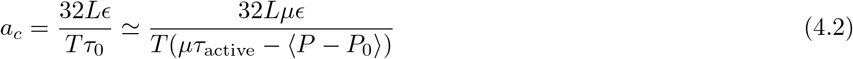

For 〈*a*〉 ≥ *a_c_*, the vein grows, otherwise it vanishes. In other words, when 〈*P*〉 is large, the vein is likely to grow, whereas it vanishes when 〈*P*〉 is small.

### 2. Parallel veins

We investigate parallel veins as shown in the network depicted in App. 4 - figure 1-D.i. The flow rate *Q* splits up into two currents in the two vein branches such that, according to Kirchhoff’s laws

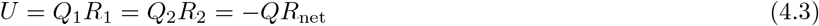

where *Q*_1_ (resp. *Q*_2_) is the flow pervading vein 1 (resp. 2). Since incoming flow rates have to sum up to zero at nodes, we also have

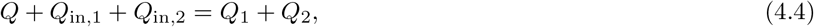

where *Q*_in,*i*_ is the net flow generated by each vein indexed by *i* over long times. The shear rates inside each of the veins are

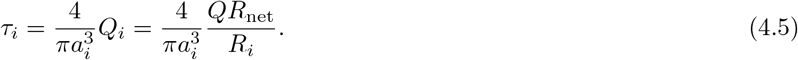

After standard calculation steps, we obtain

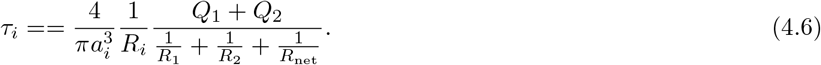

**Appendix 4 - figure 1.**
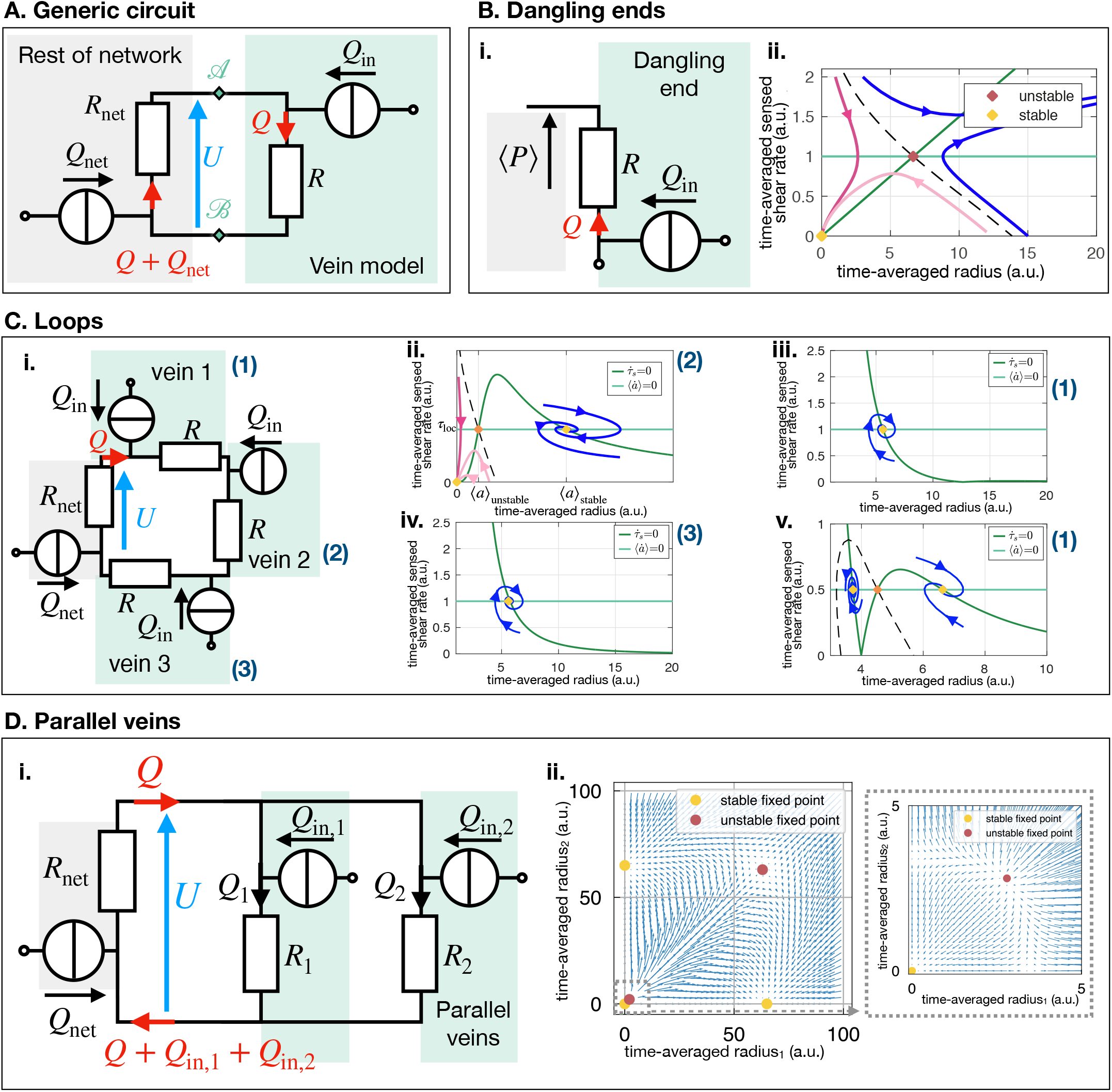
All circuits discussed in the text. (A) General circuit for a vein connected to the rest of the network. (B) Dangling ends; (i) Electric circuit and notations and (ii) stability diagram in the shear radius space. (C) Loops; (i) Electric circuit and notations (ii) Stability diagram in the shear radius space for vein 2 and (iii) for veins 1 and 3. (D) Parallel Veins; (i) Electric circuit and notations (ii) Stability diagram in the radius - radius space.

Now we remark that we can define for each of the parallel veins the resistance of the rest of the network from the viewpoint of each vein. In fact, for *R*_1_, the rest of the network is comprised of *R*_net_ and *R*_2_ in parallel. Hence from the single vein perspective of *R*_1_, we may define the equivalent resistance of the rest of the network *R*_net,1_, such that 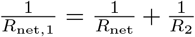. Similarly for *R*_2_. As a result we see that

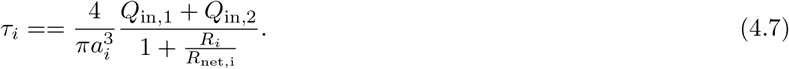

We thus coherently find that the relative resistance *R*_1_/*R*_net,1_ will determine the magnitude of *τ_i_* and hence its potential stability.

In terms of the respective vein radii, we have

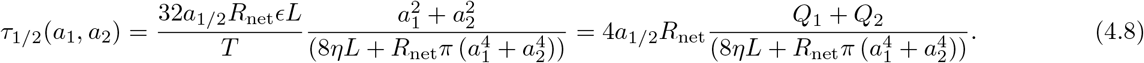

This expression allows us to draw a stability diagram in the *a*_1_, *a*_2_ space, see App. 4 - figure 1-D We find that there are three stable fixed points (0, 0), (0,*a_c_*), (*a_c_*, 0) and (*a_c_, a_c_*) is an unstable fixed point. Note that this diagram is very similar to the one obtained by [27]. As a consequence of (*a_c_, a_c_*) being unstable, one vein always shrinks in favor of the other.

We check that the instability of the parallel veins is consistent with the predictions that we could make with the resistance ratio. According to the stability diagram, one vein say of index 1 shrinks in favor of the vein with index 2 if and only if *R*_1_ > *R*_2_. In that case, we also have 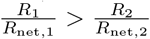. In fact we have the series of inequalities

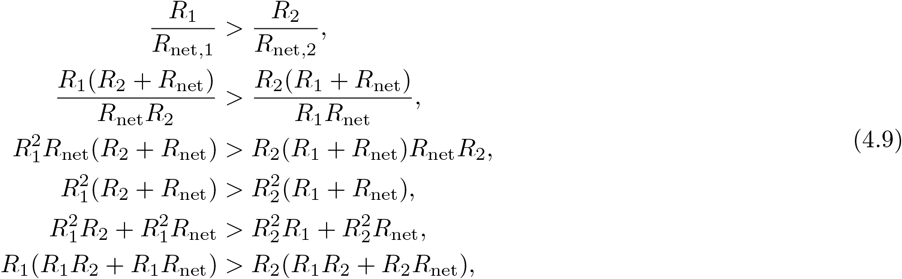

which is indeed the case since *R*_1_ > *R*_2_. Since 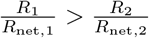, we can read off directly that vein 1 is more likely to shrink than vein 2, since the energetic gain to shrink vein 1 is bigger than that to shrink vein 2.

### 3. Loops

We investigate loops as shown in the network depicted in App. 4 - figure 1-C. Kirchhoff laws impose *Q*_net_ = –3*Q_in_* and

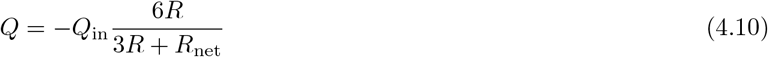

such that the shear rate through each of the veins writes as

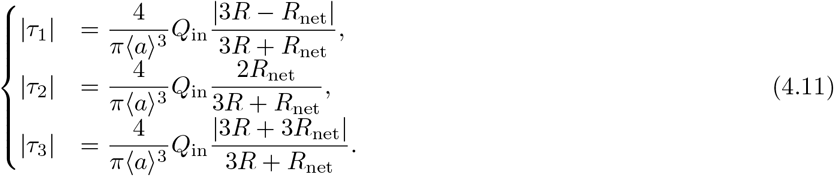

Since 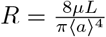, we find that

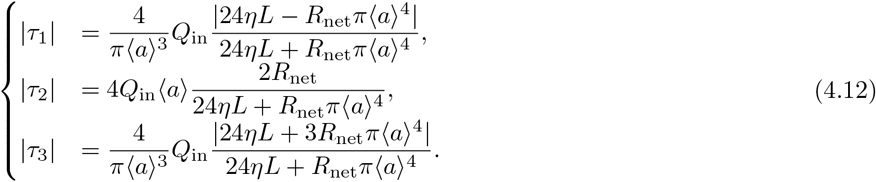

From these equations, we see that vein 2 behaves just like the generic vein (of the generic circuit). If 〈*a*〉 is small, most probably that vein will disappear - see App. 4 - figure 1-C. In general veins 1 and 3 only have one stable fixed point, and essentially have a bounded size - see App. 4 - figure 1-C (as long as 2 has not vanished yet).

